# Quantitative stable-isotope probing (qSIP) with metagenomics links microbial physiology and activity to soil moisture in Mediterranean-climate grassland ecosystems

**DOI:** 10.1101/2022.05.02.490339

**Authors:** Alex Greenlon, Ella Sieradzki, Olivier Zablocki, Benjamin J. Koch, Megan M. Foley, Jeffrey A. Kimbrel, Bruce A. Hungate, Steven J. Blazewicz, Erin E. Nuccio, Christine L. Sun, Aaron Chew, Cynthia-Jeanette Mancilla, Matthew B. Sullivan, Mary Firestone, Jennifer Pett-Ridge, Jillian F. Banfield

## Abstract

The growth and physiology of soil microorganisms, which play vital roles in biogeochemical cycling, are likely dependent on current and prior soil moisture levels. Here, we developed and applied a genome-resolved metagenomic implementation of quantitative stable isotope probing (qSIP) to an H_2_^18^O labeling experiment to determine which microbial community members, and with what capacities, are growing under *in situ* conditions. qSIP enabled measurement of taxon-specific growth because isotopic incorporation into microbial DNA requires production of new genome copies. We studied three Mediterranean grassland soils across a rainfall gradient to evaluate the hypothesis that historic precipitation levels are an important factor controlling trait selection. We used qSIP-informed genome-resolved metagenomics to resolve an active subset of soil community members and identify the ecophysiological traits that characterize them. Higher year-round precipitation levels correlated with higher activity and growth rates of flagellar motile microorganisms. In addition to bacteria that were heavily isotopically labeled, we identified abundant isotope-labeled phages suggesting phage-induced cell lysis likely contributed to necromass production at all three sites. Further, there was a positive correlation between phage activity and the activity of putative phage hosts. Contrary to our expectations, the capabilities to decompose the diverse complex carbohydrates common in soil necromass or oxidize methanol and carbon monoxide were broadly distributed across active and inactive bacteria in all three soils, implying that these traits are not highly selected for by historical precipitation.

## Introduction

Soils are among the most diverse microbial ecosystems, and microbial communities modulate the properties of soil that define its capacity to support terrestrial macro ecosystems, and human agro-ecosystems (Fierer, 2017). Microbial communities contribute to global biogeochemical cycles (Li et al., 2019; Xu et al., 2013) and soil ecosystem services (e.g. moisture retention, nutrient availability, and structure; Roberson and Firestone, 1992). Understanding microbial communities and traits along environmental gradients is foundational to predicting how soil biogeochemical processes will be altered by climate change (Cavvichioli et al., 2019). Microbial traits underlie soil carbon biogenesis (e.g.,extracellular polysaccharide synthesis) and turnover via polymer degradation and due to cell death (Sokol et al. 2022). In fact, emerging paradigms of soil organic carbon (SOC) biogenesis suggest that microbial necromass constitute much of SOC (Kallenbach et al., 2016, 2015; Liang et al., 2017). Critical gaps remain in our capacity to link microbial functional capabilities to *in situ* measurements of microbial gross growth and mortality, and also to differentiate active and inactive viral populations driving microbial community dynamics and responses to environmental variation.

Soil ecology studies frequently attempt to link surveys of microbial community composition with measurements of environmental parameters. For example, amplification and sequencing of 16S rRNA gene sequences from environmental DNA was used to show that soil pH contributes strongly to microbial community structure across diverse soils (Fierer and Jackson, 2006), and that many 16S rRNA gene-based microbial phylotypes abundant in soils are common across global soil biomes and edaphic factors (Delgado-Baquerizo et al., 2018). Sequencing phylogenetic marker genes and whole-genomes of populations of microbial cultures has also revealed the effects of latitude (Choudoir et al., 2016; Choudoir and Buckley, 2018) and soil parent material (Greenlon et al., 2019) on microbial community composition.

A limitation of phylogenetic marker-based studies is their inability to robustly predict microbial traits of actively growing taxa, i.e., the functional capacities of soil microbial communities that are relevant to the biogeochemical properties of that ecosystem’s soil. Further, marker genes such as 16S rRNA tend to be slow evolving, and so if they predict evolutionary traits they would likely be ancient. More fast-evolving traits likely define microbial niches. Genome-resolved metagenomics provides a route to access these and enables predictions of the sets of capacities of individual soil organisms without the requirement for cultivation (Butterfield, 2016; Diamond 2019), including for organisms missed in marker-gene amplicon studies (Brown et al. 2015) and viruses (Starr 2016). However, a limitation of these studies is the lack of information about which organisms are active. Although tools such as iRep (Brown et al., 2016) can use DNA sequence coverage to provide insight into replication rates, these methods have limitations (Long et al. 2020) are not very effective in soil studies due to genome completeness and coverage requirements. Metaproteomics measurements can detect abundant proteins, and thus infer bacterial activity (Butterfield et al., 2016), but the insights are often limited by extraction bias and proteins from only a small subset of the most abundant soil microbes are identified. Soil metatranscriptomics, a more encompassing and taxonomically informative analysis, may reveal how functions are expressed in space and time, e.g., that carbohydrate decomposition is conducted by distinct guilds of taxa that operate in different soil niches (Nuccio et al., 2020). However, gene expression cannot be directly linked to growth rates (Papp, 2018). Bio-orthogonal non-canonical amino-acid tagging (BONCAT) can tag cells that are actively synthesizing proteins, sort them, and sequence marker genes. This method has been applied to soils and revealed that as many as 34% of cells in soil are translationally active at any time (Couradeau, et al., 2019), but it cannot link directly to substrate usage and biosynthesis. Stable isotope probing (SIP) tracks isotopically labeled substrates into microbial populations that consume them. Thus, this method can link resource utilization to activity as a function of environmental conditions and overall community structure.

In SIP, DNA from isotopically labeled organisms is separated by density-gradient centrifugation and identified with marker gene amplification and/or metagenomics. For example, SIP experiments have been combined with metagenome sequencing to infer horizontal gene transfer events responsible for conferring isoprene degradation among novel phyllosphere taxa (Crombie et al., 2018), functional diversification between cellulose and lignin degradation in forest soils (Wilhelm et al., 2018), microbes acting in consortia for the full degradation of polycyclic aromatic hydrocarbons in contaminated soils (Thomas et al., 2019) as well as in seawater (Sieradzki et al., 2021), and linkages of uncultured microbial populations to rhizosphere carbon cycling (Starr et al., 2018). Quantitative stable isotope probing (qSIP) additionally estimates population-specific growth and death rates by tracking compositional information for 16S rRNA genes (Blazewicz et al., 2020; Koch et al., 2018). Here, we applied qSIP-informed genome resolved metagenomics, following a H_2_^18^O addition experiment, to differentiate active from inactive soil microbes, define their metabolic capacities and evaluate the potential roles of phages in bacterial cell lysis and carbon cycling among mediterranean grassland soils across a natural rainfall gradient.

## Methods

### Sample collection, isotopic enrichment and fractionation

Triplicate 0-10cm soil cores were collected from northern California annual grassland sites at Sedgwick Reserve, Hopland Research and Extension Center, and Angelo Coast Range Reserve between February and March 2018 (the period when water is most available at each site annually). Soil at these sites developed on similar parent material, are overlaid by annual grasses, including *Avena spp.,* and experience a rainfall gradient of 388 mm yr^-1^ to 2833 mm yr^-1^. Each soil core was homogenized and separated into 5 g subsamples that were dried at room temperature over 24 hours to 1.5 % gravimetric soil moisture, re-wetted to 25-30 % moisture with either natural abundance ^16^O-H_2_0 or 98.15 atom percent ^18^O-H_2_O, and then incubated for 8 days. DNA from each incubated sample was extracted and subjected to ultracentrifugation in a cesium chloride density gradient, then separated into 36 fractions (∼200 ul each) density fractions. The fractions for each sample were binned into 9 groups based on density (1.6900-1.7099 g/ml, 1.7100-1.7149 g/ml, 1.7150-1.7199 g/ml, 1.7200-1.7249 g/ml, 1.7250-1.7299 g/ml, 1.7300-1.7349 g/ml, 1.7350-1.7399 g/ml, 1.7400-1.7468 g/ml, 1.7469-1.7720 g/ml), and fractions within a binned group were combined and sequenced. Soil water retention curves were generated for Sedgwick, Angelo, and Hopland field sites using a tensiometer (HYPROP) and dew point potentiometer (WP4C) as previously described; full details of sample collection and processing are provided in Foley et al., 2022.

### DNA sequencing, assembly, annotation and binning

DNA sequencing libraries were generated using the Kapa HyperPrep kit (Roche) from each density fraction, as well as unfractionated DNA from each incubated soil sample. Paired end, 150 bp reads were generated with two lanes of the NovaSeq platform (Illumina), to an average depth of 7 gbp per library. Illumina adapter and Phix sequences were removed with BBtools (https://jgi.doe.gov/data-and-tools/bbtools/), and low-quality sequences were trimmed or discarded with sickle (Joshi and Fass, 2011).

Quality-filtered reads were assembled with Megahit (ver v1.2.9) with parameters “--k-min 21 --k-step 6 --k-max 255” (Li et al., 2015) for each individual density-fraction library as well as unfractionated-DNA libraries; co-assemblies of all sliding windows of every three adjacent density fractions (1+2+3, 2+3+4,3+4+5, etc) for each incubated soil sample; and co-assemblies of all density fractions from each incubated sample (replicate samples, i.e. cores, from the same site, were assembled and binned separately). Assemblies were filtered to remove contigs shorter than 1kb.

Contigs from each assembly were annotated using multiple sources. Open reading frames (ORFs) were predicted from assembled contigs using Prodigal v2.6.3 (Hyatt et al., 2010) with the parameters ‘-m -p meta’. We used USEARCH to identify sequences homologous to predicted ORFs in the Uniprot, Uniref90, and KEGG (Kanehisa et al., 2000) databases. We predicted 16S rRNA gene sequences using the 16SfromHMM.py script, and tRNA genes using tRNAscan-SE (Chan et al., 2019). All-fraction co-assemblies were also annotated using the METABOLIC pipeline (version 1.0) (Zhou et al., 2019).

We separated metagenomic contigs into bins representing genomes from distinct microbial populations based on sequence signatures and differential abundance across samples. Quality-filtered reads from all libraries from all samples from the same site were mapped to each all-fraction co-assembly from that site using bbmap (https://jgi.doe.gov/data-and-tools/bbtools/bb-tools-user-guide/bbmap-guide/) with the parameters “fast=t ambig=random minid=0.98“. The abundance of contigs across samples was calculated using the jgi_summarize_bam_contig_depths script from the Metabat2 software package (Kang et al., 2019). Contigs from each co-assembly were sorted into genome bins using Metabat2, Maxbin2 (Wu et al., 2016), and Concoct2 (Alneberg et al., 2014). Genome bins generated by each binning algorithm were aggregated using the Bin_refinement module of metawrap (Uritskiy, 2018). Aggregated bins from all all-fraction co-assemblies were dereplicated into representative non-redundant genomes using dRep (Olm et al., 2017) using a 99 % sequence identity threshold. The same read-mapping procedure was performed for individual-fraction assemblies, and sliding window 3-fraction co-assemblies for H_2_^18^O and H_2_^16^O incubations from soil core 2 from Hopland reserve. Metabat2 was used to generate genome bins from individual-fraction, 3-fraction, and all-fraction assemblies for these two samples, and these bins were de-replicated with dRep at 99% average nucleotide identity to determine which assembly strategy would yield the most high-quality genome bins.

We re-annotated the aggregated, de-replicated genome bins by predicting ORFs using prodigal with parameters ‘-p single’. Predicted ORFs were again annotated using USEARCH (Edgar, 2010) against Uniprot, Uniref, and KEGG, as well as METABOLIC (version 1.0, Zhou et al., 2020) and DRAM (version 1.0) (Schaffer et al., 2020). De-replicated genome bins were also assessed for the capacity for extracellular polysaccharide production through the synthase-dependent pathway by first identifying all putative secondary-metabolic biosynthetic gene clusters using the antiSMASH (v 5.0 Blin et al., 2019) pipeline with strictness set to “loose”. Putative saccharide biosynthesis clusters were further classified as synthase-dependent and by class of polysaccharide using the criteria developed by Bundalovic-Torma et al. (2019). In brief, genes from biosynthetic gene clusters that antiSMASH identified as saccharide were searched against HMM profiles for gene families from known synthase-dependent polysaccharide biosynthetic gene clusters. Clusters were classified with known polysaccharide synthesis operons if the cluster had HMM hits with e-value < 10^-5^ for three or more genes from that known operon, including for Polysaccharide synthase. We then clustered all predicted genes from all extracellular polysaccharide (EPS) biosynthesis pathways identified above into putative gene families using the protein clustering pipeline described in (Meheust et al., 2019).

We replicated an analysis from Nunan et al., 2020 correlating rRNA copy number with the number of metabolic pathways annotated “degradation” among genomes in the metacyc database. For this purpose we manually parsed the metacyc database pathways information to identify all pathways containing the term “Degradation”, analogous to Nunan et al. We identified metacyc pathways in genomes assembled from this study using MinPath (ver 1.4, Ye et al., 2009). Instead of using 16S rRNA gene copy number as a proxy for maximum growth rate, we used atom fraction excess (AFE) value.

### Virus identification and host assignment

Phage-related contigs were identified with VirSorter (Roux et al., 2015) and deepVirFinder (Ren et al., 2018). VirSorter was run in ‘virome decontamination mode, and only virus contigs identified as category 1,2,4 and 5 were kept. These viral contigs were compared to those identified by deepVirFinder, in which contigs were considered as viral if they obtained a score ≥ 0.9 and a p-value ≤ 0.05 Across all fraction assemblies, a total of 119,253 viral contigs were identified. To gain approximate ‘species-level’ taxonomic resolution (i.e., a viral population), the contigs were de-replicated through clustering at 95% average nucleotide identity (ANI) and 80% coverage (Gregory et al, 2019).

Viral populations were linked to their putative microbial hosts using a scoring approach (VirMatcher; Bolduc et al,, 2020) that was previously applied to study human gut viruses (Gregory et al., 2020). The putative microbial hosts used to establish these linkages were retrieved from 443 high-confidence metagenome-assembled genomes (MAGs) assembled in this study, and used as the host database. The different bioinformatic methods used in our scoring approach include viral matches to (a) host CRISPR-spacers, (b) integrated prophages in MAG contigs, (c) host tRNA sequences, and (d) host k-mer composition (Galiez, 2017), with the details of each method and associated scores described in (Gregory et al., 2020). The host assignments shown here only include the high- and intermediate-confidence predictions with a final score of ≥1.5.

### Calculating isotopic enrichment of microbial populations

Reads from each density-fraction metagenome library were mapped to each all-fraction co-assembly for all samples from the same site (sup. **fig. 6**). For each contig, mean coverage was multiplied by contig length and divided by the total sequencing yield of that library in order to calculate relative abundance of each assembled sequence in each density fraction (for bins, this measure of relative abundance was summed across all contigs in the bin). Relative abundances were multiplied by ng/µL of DNA recovered from ultracentrifugation for that density fraction to estimate the proportion of DNA in each density fraction belonging to each assembled sequence.

These ng/uL DNA concentrations across each incubation’s density gradient for a given sequence were used to calculate mean weighted density values for that sequence in each incubated sample. Isotopic enrichment for each bin was calculated following the methods described in Hungate et al (2015) with the following modifications. The observed GC content of each sequence was used to estimate the oxygen content of the DNA comprising the sequence, and from there the maximum mean weighted density of the sequence if all oxygens were substituted with ^18^O. Additionally, we used the concentration of DNA in each recovered density fragment to normalize the abundance (measured by read mapping) of each bin or metagenomic contig (Papp et al., 2018, Sieradzki et al., 2020)The difference between the average mean weighted density for a given assembled sequence in all three natural-abundance ^16^O replicates from that site was subtracted from the average mean weighted density in all ^18^O replicates. The proportion of this observed density shift relative to the estimated maximum density shift represents the sequence’s atom fraction excess ^18^O (AFE) (**Fig. 1C**). In reality, no organism could reach the maximum possible density shift estimated because ^18^O atom fraction excess was not 100% in the ^18^O-H_2_O incubated samples (supp table 1). This can be accounted for by calculating the fraction of maximum potential enrichment as in Foley et al., 2022. We implemented a bootstrapping procedure whereby 1000 sets of three subsamples of each set of replicates were randomly selected (with replacement), and AFE calculated from these bootstrapped samples was used to estimate confidence intervals for AFE values.

**Figure 1.**
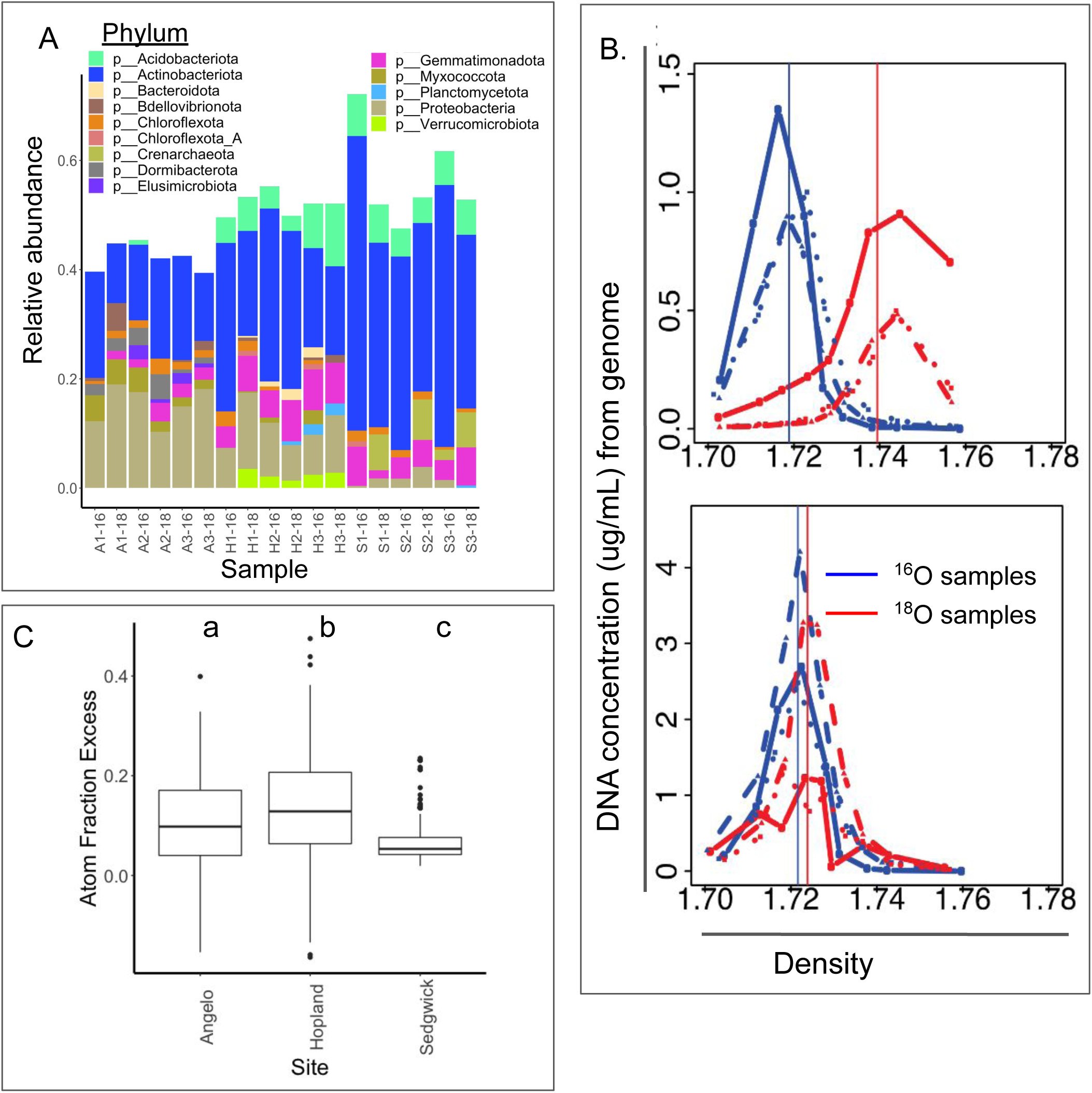
**A**: Relative abundance (from 0 to 1) and taxonomy of medium and high-quality genome bins assembled from density-gradient metagenome co-assemblies following a H_2_^18^O stable isotope probing incubation in three CA annual grassland soils (A=Angelo, H=Hopland, S=Sedgwick). **B**: Illustrative plots representing the DNA density distribution for a single organism from one site that is either active (upper panel) or inactive (lower). **C**: Boxplots showing the distribution of ^18^O atom fraction excess (AFE) values for all genome bins assembled at each site. Boxes represent the range from the 25th to 75th percentile AFE for all bins, the bold horizontal line marks the median AFE value for bins from that site. Isotopic incorporation rates across genome bins at each site are significantly statistically different from each other based on Tukey’s HSD test (indicated by the letters above boxes).

In marker-gene amplicon-based qSIP, GC is estimated based on the mean weighted density of an ASV in natural-abundance isotope samples (Koch et al., 2018 and Hungate et al., 2015). To calculate AFE on metagenomic contigs and genome bins, we used the GC content of the assembled sequence, rather than estimating GC from the mean weighted density of the sequence in ^16^O samples (we found that both methods yielded near equivalent values). In order to test the feasibility of SIP metagenomics experiments without natural-abundance samples, we further calculated AFE on each metagenomic bin using the genome’s observed GC content and observed mean weighted density in ^18^O samples (using GC to impute both maximum mean-weighted density and natural-abundance mean-weighted density) and compared these ^18^O-only AFE values to those we had calculated using ^16^O samples.

For each annotation feature (key enzymes and pathways for core metabolism, stress, complex carbohydrate degradation, physiological traits including motility, viral defense and polysaccharide biosynthesis inferred from multiple annotation packages and custom pipelines) we conducted a two-way Kolmogorov-Smirnov test comparing the AFE distribution of genomes with that trait at each site versus the AFE distribution of all genomes at the same site, correcting for multiple comparisons with the Benjamani-Hochberg procedure. We hierarchically clustered annotation features by mean AFE for genomes possessing that feature at each site (**Supp. Table 5**). The findings for the three distinct sites were then compared.

We also calculated the index of replication (iRep, Brown et al., 2016) for each de-replicated metagenomic bin by mapping reads from unfractionated DNA from each of the 18 samples to genomes co-assembled from density-fraction libraries from the same sample.

### Comparison to 16S-rRNA marker gene qSIP

Density-fractionated DNA was also used to generate 16S rRNA V4-5 variable region amplicons as described by Foley et al. 2022. Libraries were sequenced on an Illumina MiSeq instrument at Northern Arizona University’s Genetics Core Facility.

Paired-end reads were filtered to remove phiX and other contaminants with bbduk v38.56 (https://jgi.doe.gov/data-and-tools/software-tools/bbtools/bb-tools-user-guide/bbduk-guide/) and Fastq files were trimmed for quality and used to generate amplicon sequence variants (ASVs) with DADA2 v1.10 and phyloseq v1.26 (Callahan et al., 2016, McMurdie et al., 2013). Chimeric sequences were removed using removeBimeraDenovo from DADA2. ^18^O AFE of bacterial and archaeal 16S rRNA gene amplicons was quantified following a modified version of the procedure (Tag-SIP Moronado and Capone, 2016) described in Hungate et al (2015). Here, average DNA concentration was used rather than 16S-rRNA copy number (Papp, et al., 2018 and Sieradzki et al., 2020) to normalize the relative abundance of taxa within each density fraction. A weighted mean density (WMD) was then calculated for each taxon based on the distribution of its DNA across the CsCl density gradient following incubation with either natural abundance or isotopically heavy water.

Representative ASV sequences from amplicon libraries were compared to 16S rRNA gene sequences identified from metagenomic contigs using blast, filtering for any hits that aligned at 99% across the full 250 bp assembled amplicons. We calculated AFE of individual 16S rRNA gene-containing contigs using the procedure described above, and compared AFE values for 16S rRNA gene sequences calculated from amplicon sequencing and metagenomic assembly using published qSIP calculations.

## Results

### Site characteristics

To identify genomic traits associated with active microbes under varying historical precipitation patterns, we selected sites at three geographically-dispersed Mediterranean California grasslands spanning two orders of magnitude in mean annual precipitation: Sedgwick Reserve, Hopland Research and Extension Center, and Angelo Coast Range Reserve (388, 956, 2833 mm H_2_O respectively). The soils developed on similar parent material, primarily sedimentary rock including sandstone and shale, and vary only slightly in mineralogy and texture, with the driest site, Sedgwick, containing the highest proportion of clay and the highest effective cation exchange capacity (Foley et al., 2022). Sedgwick soils also reach lower water potentials at higher moisture contents than those from Hopland or Angelo (**Supp. Fig. 1**). All three sites had similar nitrogen and carbon content (**Supp. Table 1**; Foley et al., 2021).

### Density fractionation effects on genome recovery

To evaluate the effect of density fractionation and isotope labeling on metagenome assembly, we assembled and binned individual fractions, sliding windows of co-assemblies of 3 adjacent fractions, and co-assemblies of full density gradients.

For one sample (Hopland replicate soil core 1, both ^18^O and ^16^O incubations), we compared genome recovery outcomes using DNA sequences from all fractions vs. three adjacent fractions on the density gradient and found that the all-fraction co-assemblies yielded the largest number of high-quality genome bins (**Supp. fig. 3**). Where genomes were recovered by both approaches, the highest quality bins (defined by maximum completeness and minimum contamination) tended to be those from the full coassembly and most good bins that were recovered only once came from the all-fraction co-assemblies. We clustered genome bins from each assembly at 99% sequence identity, and found that of 65 genomes representative of the clusters, 44 with the highest quality (based on dRep scores) were from the all-fraction assembly. Twenty-eight clusters contained MAGs recovered in multiple cross-fraction co-assemblies, in 15 of which, the highest quality genome was from the all-fraction assembly (**Supp. fig. 3**). Among the 37 clusters that we recovered in only one co-assembly, 29 were assembled in the all-fraction co-assembly (**Supp. fig. 4**). Read mapping indicated that genome bins had coverage across the density gradient, consistent with the additive effects of combining density-fraction libraries contributing to improved assemblies (**Supp. Fig. 5**). In addition, relative coverage across the density gradient varied systematically (with low GC regions having relatively higher coverage at lower densities). Based on the superior performance of the all-fraction co-assemblies, we proceeded with these assemblies for each sample for genome binning, annotation, isotopic-label quantification, and statistical analyses.

### Metagenome assembly and binning

From the all-fraction co-assemblies from all samples, we reconstructed 433 non-redundant genome bins with estimated completeness > 75% and contamination < 25%, representing a diverse array of common soil associated microbial taxa, with Actinobacteria predominating at all three sites (**Fig 1A**). Diverse Proteobacteria are abundant at Angelo and Hopland reserves. Less-abundant organisms diverge between sites: we detect Gemmatimonadetes only at Angelo; Bacteroidetes and Chloroflexi predominantly at Hopland; and Chrenarchaeota only at Sedgwick. *Bdellovibrio* was observed in Angelo and Hopland soils.

### Quantifying isotopic incorporation in metagenomic sequences

To associate microbial ecophysiological traits with metabolic activity and population turnover, we calculated atom fraction excess (AFE) of ^18^O in DNA sequences assembled from qSIP metagenomes (**Fig 1B** **and Supp. Fig 5**). Isotopic enrichment represents new incorporation of oxygen into biomass, and AFE is therefore proportional to metabolic activity and population growth of the organisms from which the sequence assembled (Blazewicz et al., 2011). Estimated AFE ranged from -0.16 (reflecting experimental error) to 0.47, with the range and distribution of AFE varying significantly by site (**Fig 1C**, **Supp. Table 2 and 3**). We refer to organisms with AFE lower than average for the site where it was assembled as ‘low activity’ and those with higher than average AFE as ‘high activity’. In the dataset of ^18^O-SIP 16S rRNA gene amplicon sequences that parallel our ^18^O-SIP metagenomic libraries, we found a significant positive correlation between activity estimates from the two data types (**Supp. Fig 7**).

The index of replication (iRep) provides an orthogonal measure of *in situ* microbial growth calculated from metagenomic sequence data (Brown et al. 2016). However, only 3 bins exceeded the coverage filtering threshold required for iRep. Of these three bins, iRep was inversely related to AFE calculated for each (**Supp. Table 4**), suggesting the difference in measuring instantaneous replication rates (as in iRep) versus gross population growth over time (qSIP).

AFE from qSIP expresses the observed shift in density as a proportion of the maximum theoretical density shift for an organism’s genome. This shift is calculated by subtracting an organism’s density in unlabeled samples (determined by its GC content), from its density in labeled samples. However, if the observed density of an organism’s unlabeled genome matches the theoretical density calculated from the genome’s GC values, we should be able to infer an organism’s isotopic incorporation purely from the observed density of its isotopically-labeled genome. With our dataset, we calculated AFE for each genome bin using only read coverage data from ^18^O-enriched libraries and found a very strong and significant positive correlation between AFE values calculated from ^18^O and ^16^O libraries (p<2.2×10^-16^, R^2^=0.93, **Supp. Fig. 8**). Given this result, it is possible that unlabeled samples need not be sequenced in future SIP-metagenomics studies so long as substantial genome recovery is possible.

We found ^18^O AFE varied widely for genomes within the same phylum, both within and between sites (**Fig 2A**), with community-level AFE distributions reflecting measured respiration rates between sites (**Supp. Fig. 2**). Actinobacteria, the most abundant phylum observed at all three sites, had a particularly wide range in activity, including the highest and lowest AFE values at each site. Most Actinobacteria genomes had similar AFE distributions relative to the broader microbial community at each site, while members of the family Nocardioidaceae had higher activity levels than other Actinobacteria at all three sites.

**Figure 2.**
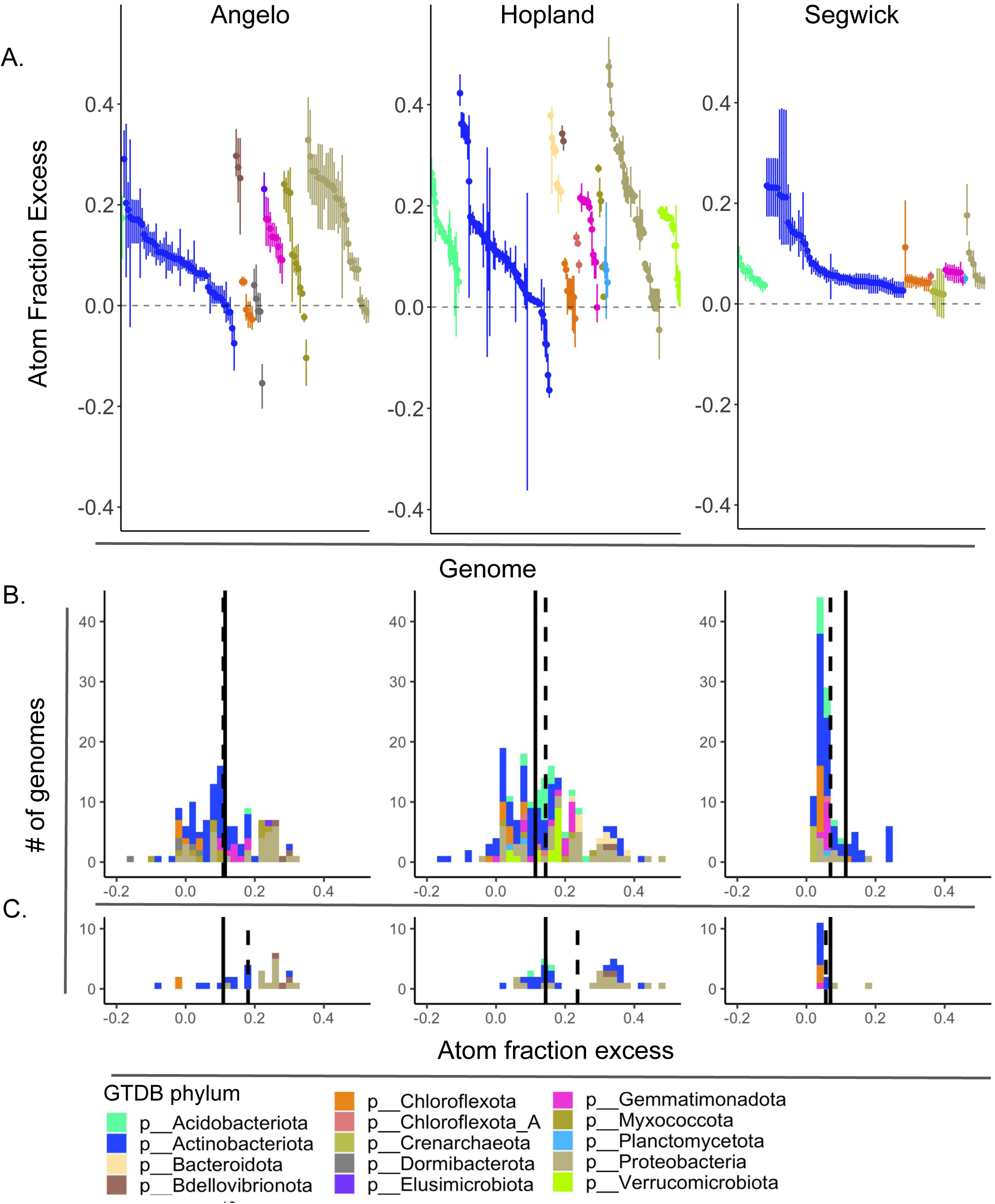
**A**: Range of ^18^O atom fraction excess (AFE) confidence intervals for all genome bins at each of three CA grassland sites colored by phylum **B**: Distributions of AFE values for all bins at each site. Solid line represents the average AFE across all bins at all sites, dotted line represents average AFE across bins from the site shown in that figure panel. **C**: Distribution of AFE values for genome bins encoding flagellin at each site. These genomes are significantly enriched at Angelo and Hopland relative to average bins at each site. Solid line represents the average AFE from all bins at that site; dotted line is the average AFE of genomes encoding flagellin genes at that site.

Members of several less common groups of bacteria had more consistent activity levels compared to Actinobacteria. Chloroflexi and related phyla (Chloroflexota_A, also known as Rif_CHLX, and bacteria of the phylum Dormibacterota) at all three sites and Chrenarchaeota from Sedgwick had consistently very low activities. Members of the phylum Planctomycota had low activity levels (AFE < 0.1) at both Hopland and Sedgewick sites.

*Bdellovibrio* (known as “predatory” bacteria for their obligate intracellular parasitism of other bacteria), represented by four genomes from Angelo and Hopland, were among the most active organisms at each site. Consistent with obligate parasitism, the genomes all contain loci for type IV pilus known to be involved in host attachment, and exhibit many auxotrophies in amino-acid biosynthesis.

### Statistical testing distinguishes metabolisms and ecophysiological traits by activity across sites

To identify microbial traits associated with activity and how these patterns were framed by our soils’ different precipitation regimes, we used a statistical test to identify traits where the AFE significantly increased with higher site precipitation (**Fig. 2B**).

Motility emerged as a strong predictor of growth. Putatively motile organisms (genome bins encoding full or mostly full suites of genes for flagellar biosynthesis) were statistically overrepresented among active organisms from the wetter Angelo and Hopland sites (organisms with flagella-encoding genes had AFE 8.1 and 9.2 higher, and 1.3 % lower than all organisms assembled at Angelo, Hopland, and Segwick respectively, **F**i**g. 2C, 3A**). This is partially due to the presence of flagellar genes in the highly-active Bdellovibrionata at Angelo and Hopland. Similarly, active Nocardioideaceae (phylum: Actinobacteriota) at Angelo and Hopland have the capacity for flagellar motility whereas those from Sedgwick were not.

**Figure 3.**
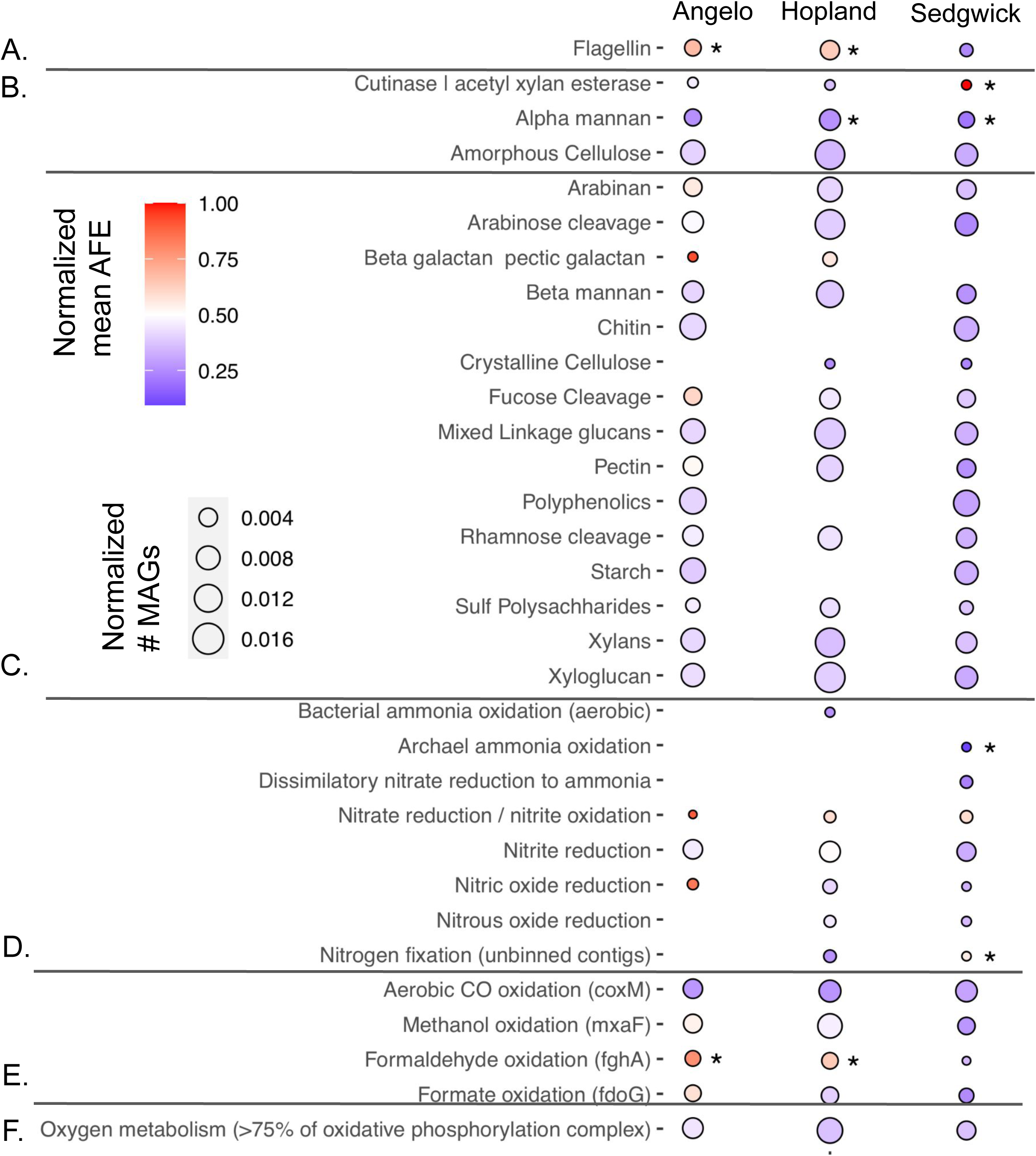
Heatmap of number of genomes with annotated functions and the activity (reflected by ^18^O atom fraction excess-AFE) of those genomes, at three grassland sites in northern CA, for genomes encoding: **A**: flagellin (significantly active at high moisture sites. **B**: cutinase (significantly active at low moisture site, Sedgwick) **C**: Polysaccharide metabolism **D**: C1 metabolism **E**: Nitrogen metabolism **F**: Oxygen metabolism. For all panels, the size of each point is proportional to the number of genomes with that trait. Circle color represents the average AFE of genomes with that trait, per site. An asterisk next to a point indicates significant difference from the average AFE of all bins at that site.

We hypothesized that organisms with limited mobility would tend to be more versatile in terms of the substrates that they can metabolize. We found that the non-motile Nocardioideacae from Sedgwick encoded higher numbers of polysaccharide degradation pathways. These organisms also had higher numbers of enzymes for nitrate reduction and nitrite reduction, whereas organisms from Angelo and Hopland only had the capacity for nitrite reduction.

Genes for carbon monoxide and methanol dehydrogenases were broadly distributed phylogenetically, geographically, and across activity levels measured by AFE (**Fig. 3E**). Carbon monoxide dehydrogenase (coxL) was primarily confined to Actinobacteria across sites (**Supp. Fig. 9**). Calcium-dependent methanol dehydrogenase (mxaF) was broadly distributed among Acidobacteria,

Gemmatimonadetes, and Proteobacteria (**Supp. Fig. 10**). Genomes annotated with mxaF were often highly active at Angelo (0.151 for mxaF bins, average AFE of 0.107 for all Angelo bins), and to a lesser extent at Hopland (0.174 compared to 0.135) but bacteria with mxaF from Segwick were relatively inactive (0.063 vs 0.110).

We found significant positive correlation between annotated metacyc degradation pathways and AFE-based activity levels at Angelo, but not the other soils (**Supp. Fig. 11**). Across sites, we aggregated genomes based on phylum affiliation and found no significant correlation between substrate diversity and activity levels. We also examined the distribution of degradation pathways for several complex carbohydrates and found that of genomes possessing any of 13 polysaccharide degradation pathways curated in the DRAM genome annotation pipeline (Schaffer et al., 2020), none differed significantly in inferred activity levels from the total community represented by the assembled metagenome at any site (**Fig. 3C**). We observed significant correlations between the diversity of polysaccharide degradation pathways encoded in a genome and the high activity levels for that genome in some phyla, but the phyla that displayed this pattern often varied across sites. For example, we found a significant positive correlation between the number of polysaccharide-degradation pathways encoded in Proteobacteria genomes and organism activity (AFE) across all sites (r = 0.47, p<0.05) whereas for Gemmatimonadetes, there was only a significant correlation between polysaccharide-degradation pathways and activity at Angelo (r=0.33,p<0.05) (Supp. **Fig. 12**). Conversely Chrenarchaeota at Sedgwick and Acidobacteria and Bacteroidetes from Hopland exhibited higher activity correlated with a lower diversity of polysaccharide degradation pathways.

MAGs from the three sites varied in the presence of specific nitrogen cycle enzymes, in the abundance of nitrogen-related genes, and in the inferred activity levels of bacteria involved in nitrogen compound transformations (**Fig. 3D**). All three sites had organisms whose genomes encoded the first three steps of denitrification (reduction of nitrate to nitrite; nitrite to nitric oxide; nitric oxide to nitrous oxide). The capacity for nitrate reduction was only observed in genomes of highly active bacteria from the wettest site, Angelo, whereas genes for nitrite reduction were broadly distributed among organisms with varied activity levels at all three sites. Nitric oxide reduction capacity was also encoded in genomes of the most highly active bacteria from the Angelo site.

The capacity for nitrous oxide reduction to molecular nitrogen was only observed in genome bins at Hopland and Sedgwick (organisms potentially capable of N_2_O-reduction were close to average activity at both sites). The capacity for aerobic ammonia oxidation was predicted for Acidobacteria from Hopland, and Crenarchaeota from Sedgwick, but the inferred activity levels of these organisms were low. In addition, genes for nitrogenase were encoded on unbinned contigs from the Hopland and Sedwick datasets. A few nitrogenase operons were encoded on high AFE contigs from Sedgwick.

We annotated genome bins for EPS biosynthetic gene clusters for a class of polysaccharides produced through the synthase-dependent biosynthetic system including poly-N-acetylglucosamine (PNAG), cellulose and acetylated cellulose, alginate, and Pel. PNAG was the most commonly-observed polysaccharide biosynthesis cluster, assembling in a broad range of taxa with varied activity levels at all sites (**Supp. Fig. 13**). Notably, Actinobacteria of the family Nocardioideaceae –that were apparently highly active at each site– are not predicted to synthesize PNAG. *Quantifying activity in phage:*

Many studies have documented viruses in soil (e.g. Williamson et al., 2017, Trubl et al., 2021, ter Horst, et al., 2021), but as viruses can be deactivated in soil through multiple mechanisms (e.g., by sorption to minerals), a grand challenge is to assess what fraction of viruses is active. We identified potential phage sequences in metagenomic assemblies (total of 119,253 viral contigs across all all-fraction assemblies; clustered into 8,617 viral populations), and examined patterns of activity measured by AFE, focusing on phage contigs for which we confidently predicted hosts. As with microbial genomes, putative phage contigs demonstrated a range of AFE at each site. We find a significant positive relationship between the AFE of a host genome and its putative phage genome (p-value < 2.45e-14, R^2^ = 0.683, Fig 4). For example, we identified a circular 38 kbp genome for a phage predicted to infect a *Bdellovibrio* at Hopland. Circularization indicates that the sequences were derived from phage particles, not from prophage. Both the putative *Bdellovibrio* phage and the only *Bdellovibrio* genome assembled from Hopland are predicted to have been highly active (AFE ∼ 0.3-0.35 range). However, notably, for this phage-host pair and in general, hosts predicted to be highly active had lower relative abundances. High activity of hosts represents high growth rates, and we might expect that this would lead to high abundances. However, it is possible that the most active phages had infected and lysed their hosts (leading to low abundance) before samples were collected. Alternatively, the phages may simply have infected low abundance yet active hosts.

**Figure 4.**
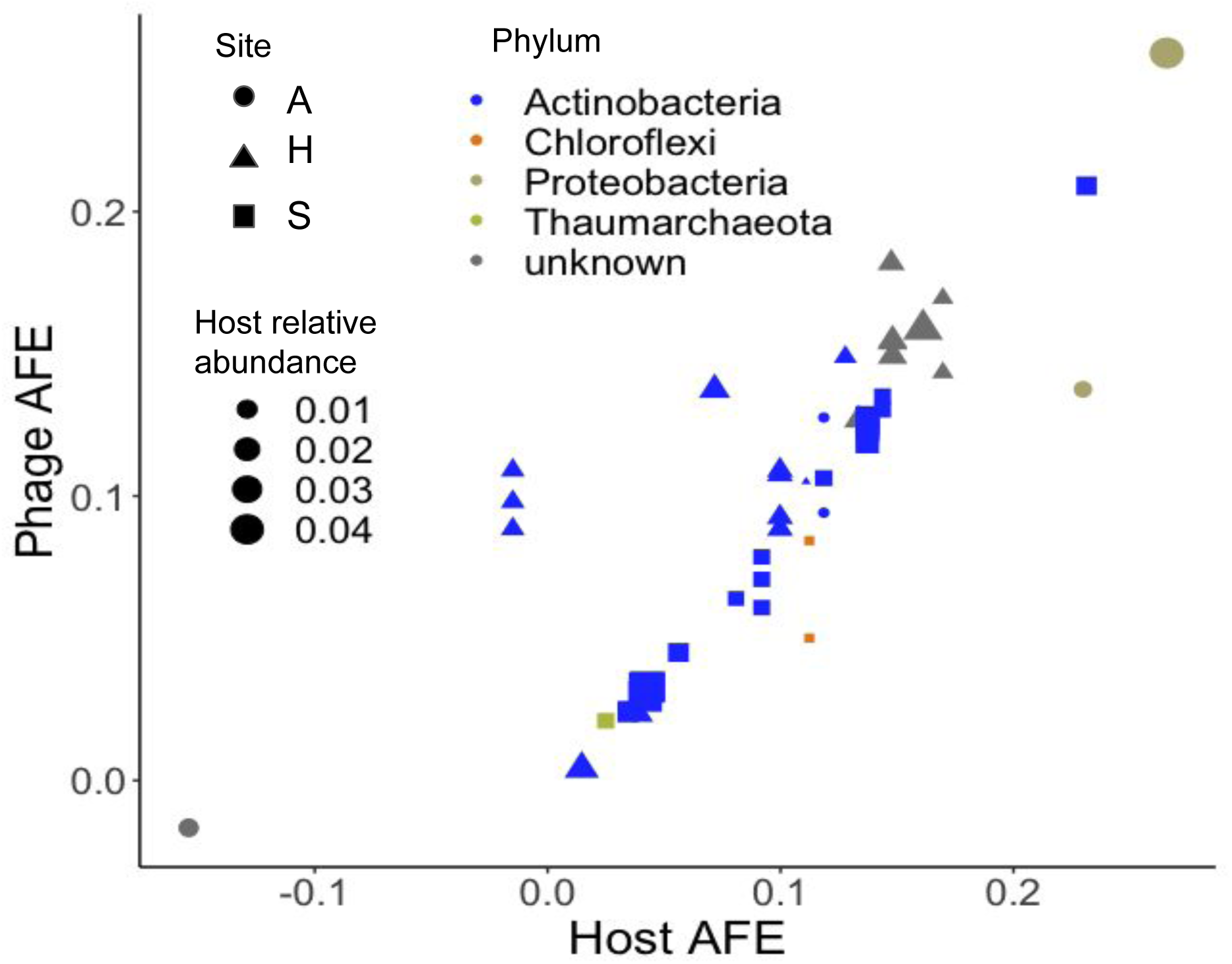
Taxon-specific activity (measured by ^18^O AFE) of grassland soil bacterial genomes versus phage predicted to infect each bacterial host based on matching CRISPR spacer sequences. Sites include Angelo (A), Hopland (H), and Sedgwick (S), which exist along a rainfall gradient in Northern California. Host relative abundance is indicated by the size of datapoints.

From all three sites, we reconstructed and annotated seven genomes of phages that encode alginate lyase carbohydrate-active enzymes (CAZYmes). Alginates are polysaccharides that may mask receptor sites used by phages during infections, and the phage lyase may circumvent this defense. When aligned to the refseq database, genes within these putative phage contigs that align to annotated genes from known organisms have highest homology to sequences from Pseudonocardiaceae, consistent with these phage infecting Pseudonocardiaceae cells. Interestingly, only five Pseudonocardiaceae from Sedgwick have operons implicated in alginate synthesis (**Supp. Fig. 14**). Putative Pseudonocardiaceae-infecting phages from Segwick that encode alginate lysase occur on metagenomic contigs that are not circularized. These contigs have higher AFE than the predicted alginate-producing Pseudonocardiaceae host genomes assembled from Segwick, consistent with their existence as phage particles that replicated in an actively growing host (**Supp. Fig. 15**).

## Discussion

Using qSIP-informed genome-resolved metagenomics we quantified the activities of bacteria, archaea and phages from the three annual grasslands that exist along a strong precipitation gradient. Previous SIP metagenome studies have typically sequenced only the heavy fraction of DNA density gradients for labeled and unlabeled samples. At best, this approach can make a binary distinction between organisms that have or have not incorporated the stable isotope label, and many organisms that incorporate the isotope are missed due to lower yet significant isotope incorporation.

Additionally, regions of the same genome with differences in GC content distribute across the density gradient, reducing assembly quality if only subsets of the density gradient are assembled. Finally, as the mean weighted density is determined by both GC content and isotope incorporation, many low GC genomes with high AFE may be missed if only the heavy fraction was sequenced. By sequencing across the density gradient, the approach used here, we recovered more and higher quality genomes and quantified isotopic enrichment in microbial and phage genomes. Estimates of AFE for metagenomic contigs and genomes containing 16S rRNA genes align with AFE estimates for identical 16S rRNA gene sequences calculated from the paired dataset from the same samples. This approach allowed us to deduce quantitative relationships, to test for a correlation between abundance and growth, and to use a statistical framework to associate microbial traits with activity as a function of historical water inputs.

The strongest statistical signal we saw relating a microbial trait to differences across the annual precipitation gradient represented by our sample sites was flagellar motility. Genomes predicted to belong to motile organisms were proportionally most active at the wettest site (Angelo), followed by the more moderate precipitation site (Hopland), whereas the seasonally driest site (Sedgwick) had motile bacteria with lowest activity under our experimental conditions. Previous modeling has predicted that spatial heterogeneity of soil selects for organisms capable of rapidly responding to changes in availability of a wide diversity of substrates (and against alternative nutrient acquisition strategies such as chemotactic motility)(Nunan et al, 2020), whereas we found selective pressure favored increased motility in soils with a historical pattern of higher soil moisture availability.

The higher soil moisture levels at Angelo and Hopland could make motility advantageous (except for those bacteria that rely on soluble organic compounds, as diffusion rates would increase with increased soil moisture). Interestingly, the motile bacteria from family Nocardioidaceae at Angelo and Hopland have relatively few enzymes for degradation of insoluble carbohydrate compounds such as arabinan, and xylan, as well as polysaccharides containing fucose or rhamnose, compared to non-motile bacteria from this family at Sedgwick. Thus, the Actinobacteria at higher-moisture sites appear to be relatively specialized (from the perspective of carbohydrate degradation) compared to those from Sedgwick. These findings support the premise of some ecological models (Nunan et al, 2020) that suggest homogeneity associated with wetter soils should select for organisms with relatively specialized metabolisms whereas heterogeneous environments such as drier soils should select for versatile heterotrophs. In wetter soils, microbes that are motile can relocate to sites where resources they can use are found whereas in dry soils, motility is restricted so it is beneficial to be a versatile heterotroph. During the soil incubations, water potential varied little between sites. The observed patterns in microbial activity therefore reflect the effects of historical differences between sites in soil moisture. Long term climate shifts towards decreased precipitation might select for less motile organisms and their associated lifestyles.

The higher activity of motile organisms in wetter soils was driven in part by the activity of motile *Bdellovibrio* (representing 20% of flagella-encoding genomes at Angelo and 9.7% at Hopland). *Bdellovibrio* are often intracellular parasites of other bacteria. If the *Bdellovibrio* in the soils studied here are also parasites, then it may be reasonable to hypothesize that motility enabled by higher moisture levels enhanced the success of parasitic bacteria. There is increasing evidence for the predominance of microbial necromass in SOC (Kallenbach, 2016), therefore we speculate that *Bdellovibrio*-induced lysis could contribute substantially to SOC pools in wetter soils (Hungate et al, 2021).

The lack of difference in SOC abundances in the three soils may be due to increased access to the lysate as a C source in wetter soils. Climate change driven reduction in moisture levels may select against *Bdellovibrio,* lowering the contribution of bacterial predation to cell death.

Our SIP-metagenome data show that despite the precipitation gradient amongst our sites, metabolic capacities such as aerobic respiration, CO oxidation and methanol oxidation were linked to genomes whose isotopic enrichment values fell close to average community enrichment values at all three sites. Thus, we conclude that these are widespread capacities in grasslands during seasonally high soil moisture.

Metagenomic qSIP enabled us to track activity of phage populations. The observation that AFE of phage genomes is closely linked to the AFE of their predicted microbial hosts might suggest that phage lysis rates are sometimes predicted by microbial growth rates. However, a subset of these may have been prophage and were isotopically labeled during host replication. The observed low relative abundances of host bacteria in soil may be a consequence of phage replication leading to host bacterial death, but other explanations (including bacterial and eukaryotic predation) likely also contribute. It is interesting to note that phage predation could have reduced to undetectable levels the abundances of bacteria that were recently highly active, precluding deduction of their recent high activity via qSIP-informed genome-resolved metagenomics. The detection of phages with high ^18^O values but no apparent host thus may be an indication of prior host growth and a useful signature of high rates of phage-induced mortality, as well as an indicator of top-down trophic influence in soil microbiomes (Hungate et al., 2021).

The fast replication of organisms encoding enzymes involved in nitrate, nitrite, and nitric oxide reduction suggests denitrification to N_2_O likely occurs in Angelo soils during the wet season. The genomic capacity of active organisms for nitrous oxide reduction, leading to full denitrification to dinitrogen gas, was more prevalent at Hopland and Sedgwick compared to Angelo. Given that all three sites have similar nitrogen compound concentrations, we suggest that activity differences might have occurred because the sites weren’t equally wet at the time we sampled (Sup. Fig. 1A). Thus, long-term differences in yearly moisture availability likely strongly impact microbial transformations in soil nitrogen cycles. Although the bacteria capable of reduction of nitrous oxide were not particularly active at the time of sampling, these bacteria could limit emissions of N_2_O resulting from partial denitrification from the drier soils under other conditions. More importantly perhaps, the ability to reduce N_2_O to N_2_ is most prevalent in bacteria from both Sedgwick and Hopland that are non-motile. Thus, future decreases in soil moisture levels could lead to decreased emissions of this greenhouse gas. Conversely, the lack of genomes encoding genes for reduction of N_2_O at the Angelo site may predict a higher capacity for release of this greenhouse gas from Angelo compared to other soils, especially under wetter conditions. This is consistent with previous observations of higher N_2_O fluxes from wetter soils (Chen, Mothapo, and Shi, 2014).

### Methodological advances and implications

Past metagenomic SIP studies have claimed that either the density fractionation or stable-isotope labeling itself improves metagenomic assembly by reducing the complexity of the microbial community sampled within each individual fraction. In contrast, in our study we observed that co-assembling all fractions from the same sample almost always improved assembly quality, probably because it increased the coverage per genome. This suggests a possible SIP metagenomics experimental design wherein assemblies primarily come from unfractionated DNA sequenced to high depth and density fractions are sequenced at low coverage for identifying bins through differential abundance and for estimating isotopic incorporation.

In the future, this genome-resolved approach could be greatly expanded by using substrates labeled with ^13^C (e.g. Zwetsloot et al, 2021) or ^15^N labeled (e.g. Wilhelm et al., 2021, Morrisey et al., 2018), as demonstrated by 16S rRNA gene amplicon qSIP studies. Use of ^13^C and ^15^N labeled substrates could be used for in depth analysis of soil carbon biogeochemistry, because compounds predicted from genomes to be consumed or synthesized by isotopically-enriched bacteria can be quantified with spectroscopic techniques that can differentiate isotopically labeled molecules (e.g. HPLC, NMR). Further, sequencing unfractionated metagenomic DNA from not one (as was done here) but many time series points would constrain the relative abundances of specific bacteria, potentially enabling direct estimates of microbial net growth (i.e. birth and death) rates (Koch et al., 2018, Blazewicz et al., 2020). This would complement measurements of isotope incorporation into phage populations that are predicted to infect specific microbial populations, allowing linkage of cell death to phage predation.

Genome-resolved SIP can be cost and labor intensive. Given that the mean-weighted densities for organisms in natural-abundance isotope treated samples are close to those estimated purely from GC of the same sequence assembled in heavy-isotope labeled samples, it may be possible to conduct future SIP experiments with few or no natural-abundance isotope controls. This would double, for the same cost, the number of samples that could be included in metagenomic qSIP experiments, enabling deeper replication coupled with increased statistical power (Sieradzki et al., 2020) and/or more diverse treatments.

### Conclusions

The qSIP-enabled implementation of genome-resolved metagenomics that involved sequencing multiple density fractions enabled us to probe the activities and functional potential of bacteria and phages in soil microbial communities. We differentiate phenomena associated with soils that have experienced a gradient of annual precipitation levels and identify traits that may change in prevalence as climate changes, including a subset that could impact soil greenhouse gas emissions.

## Acknowledgements

We thank QB3 for sequencing support, and Rohan Sachdeva and Shufei Lei for bioinformatics support. This research was supported by the U.S. Department of Energy, Office of Biological and Environmental Research, Genomic Science Program ‘Microbes Persist’ Scientific Focus Area (#SCW1632) at Lawrence Livermore National Laboratory (LLNL) and subcontracts to Northern Arizona University and Ohio State University and Ohio State University. Work conducted at LLNL was conducted under the auspices of the US Department of Energy under Contract DE-AC52-07NA27344. The Innovative Genomics Institute-based computational facility and the Ohio SuperComputer is acknowledged for computational support.

## Data availability

Sequence data generated for this manuscript is available at NCBI under Bioproject PRJNA718849. Microbial genomes bins and annotations are available at: https://ggkbase.berkeley.edu/wsip-metawrap-drep-bins/organisms. Scripts for the computation and analysis used to generate results for this manuscript will be available at https://github.com/alexgreenlon/wsip/tree/master.

