## Supplemental figures for "Quantitative stable-isotope probing (qSIP) with metagenomics links microbial physiology and activity to soil moisture in Mediterranean-climate grassland ecosystems"

Supplemental figure 1

Figure 1
A

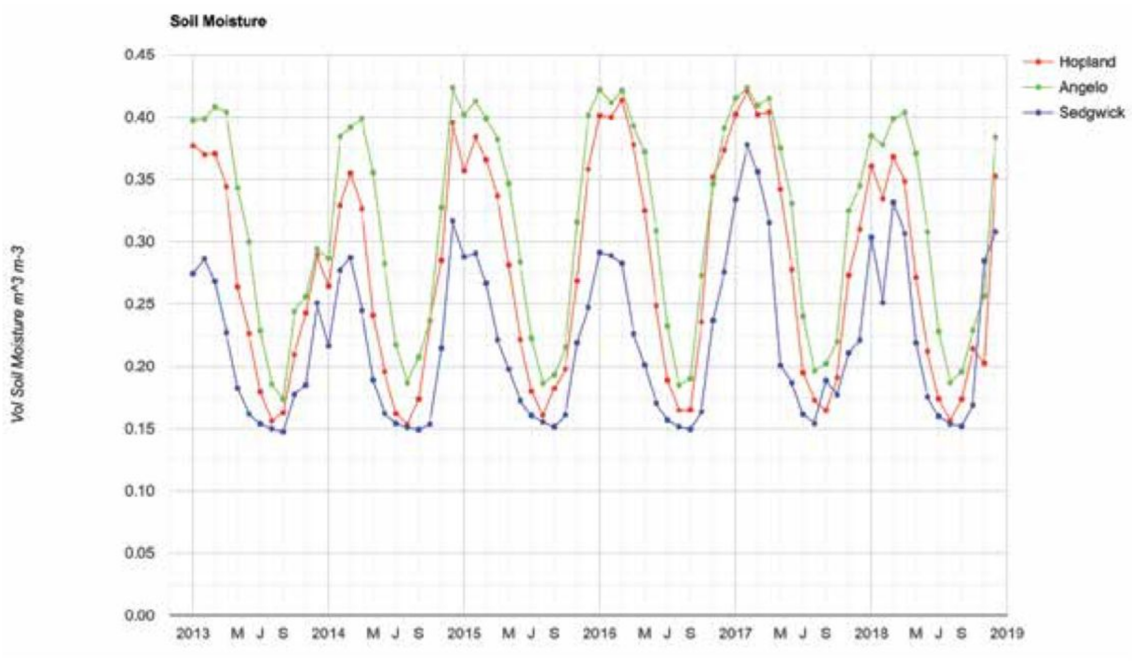


B

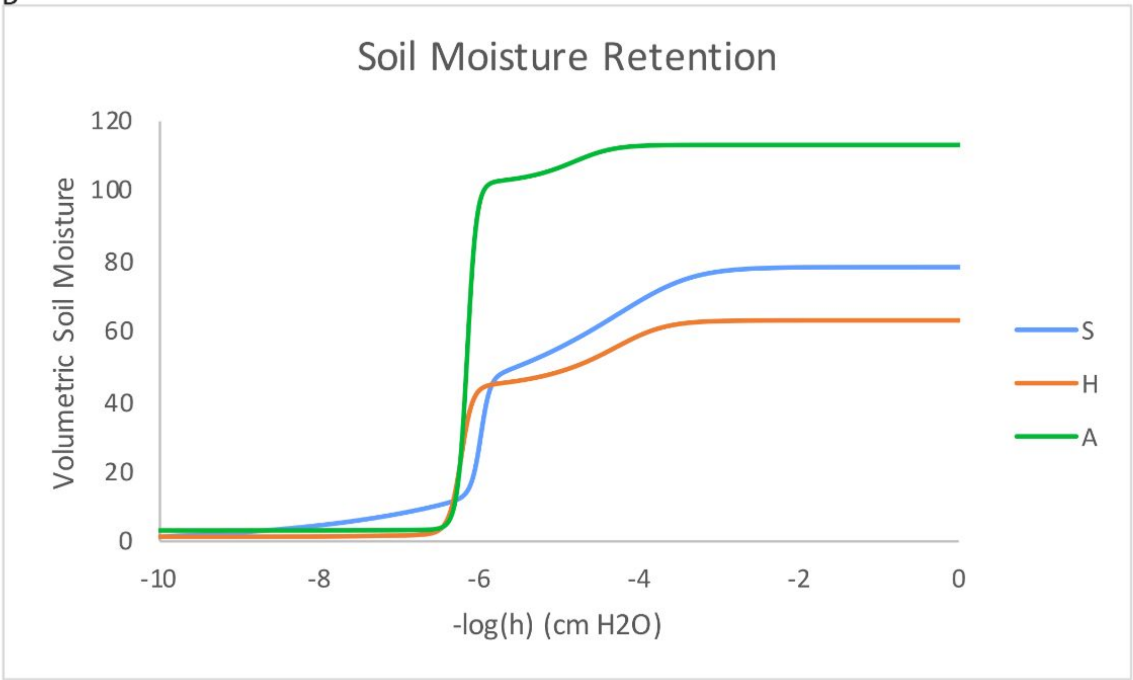


A: Yearly precipitation patterns at all three sites.
B: Moisture release curve for soils from all three sites.

Supplemental Figure 2

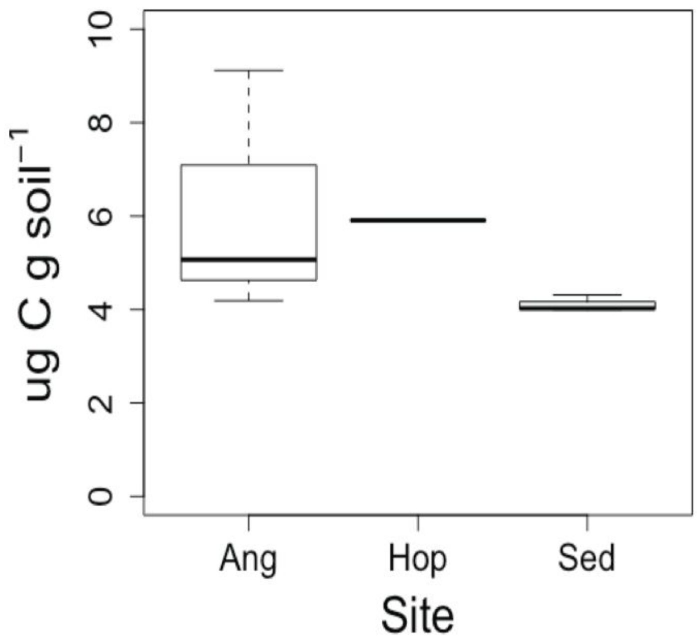


Total respiration during the incubation expressed as ug C from CO2 respired per g of soil.

Supplemental Figure 3

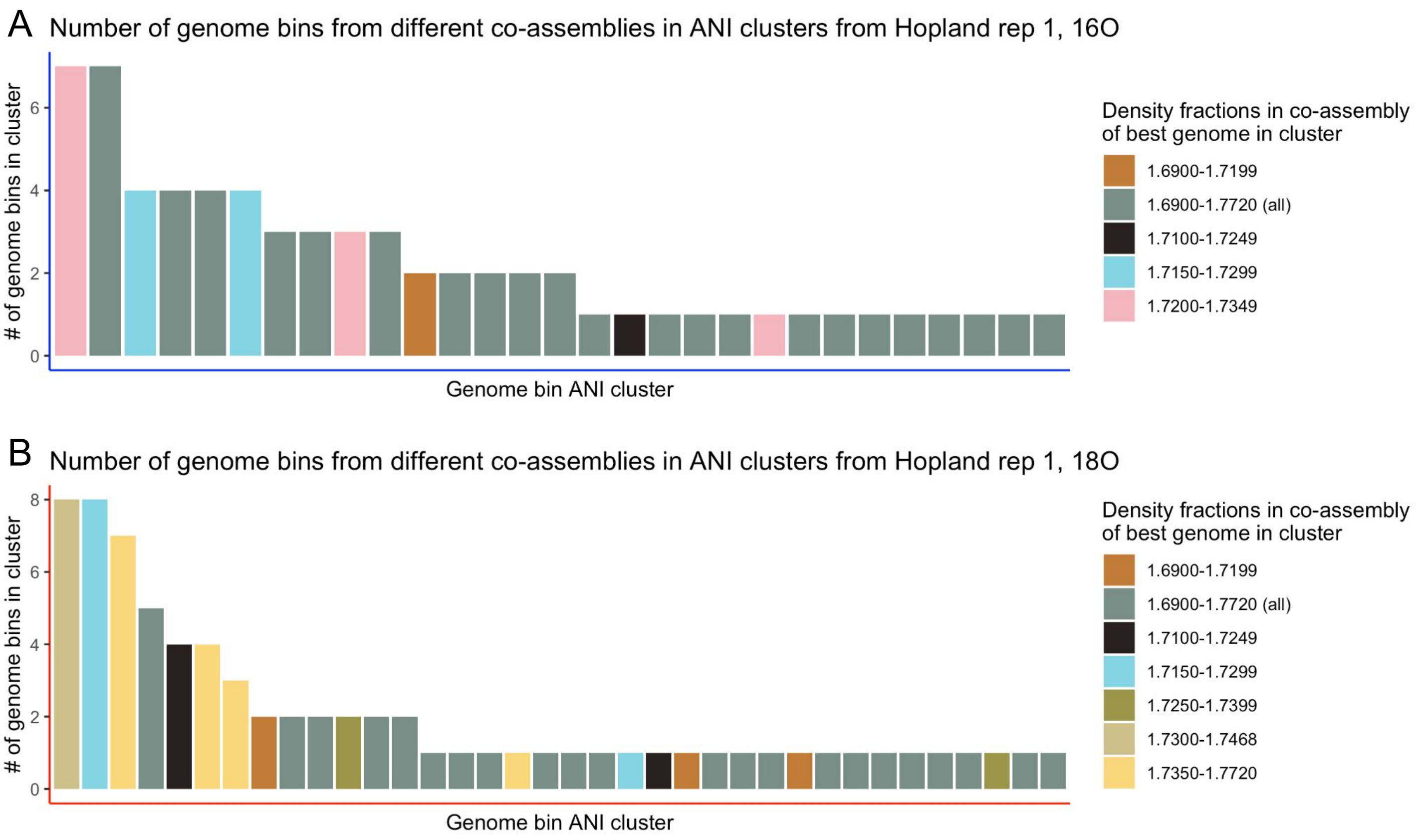


Each bar represents a cluster of identical genomes assembled in all-fraction co-assemblies as well co-assemblies of sliding windows of 3 adjacent fractions. The genome clusters were generated using dRep at 99% ANI. The height of the bar represents the number of different co-assemblies in which the genome assembled. The color of the bar corresponds to the co-assembly in which the highest quality genome assembled. For many of the most frequently assembled genomes, the highest quality genomes came from a 3-fraction co-assembly, but there were many genomes that only assembled in the all-fraction co-assembly.

### Supplemental Figure 4

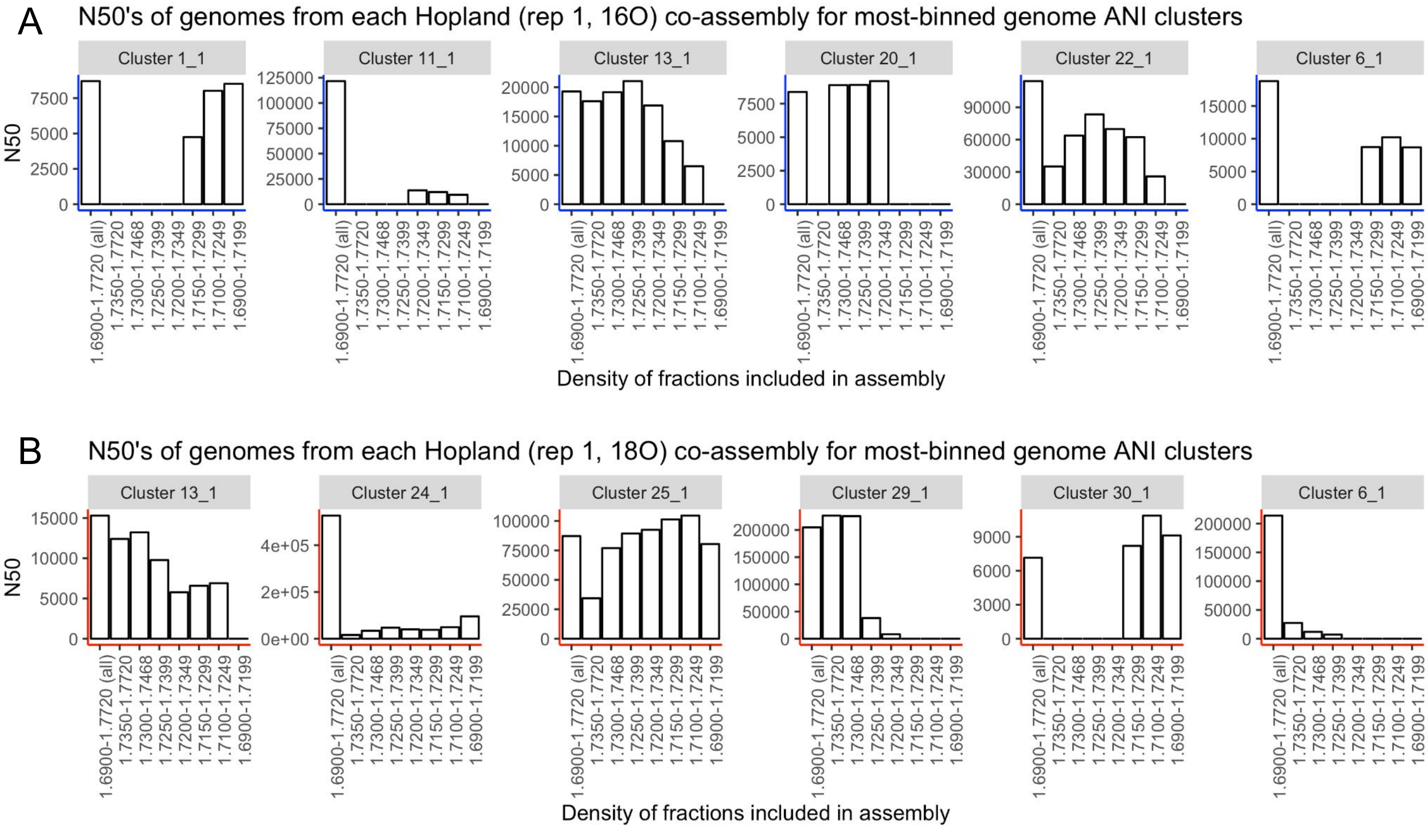


Each graph summarizes genome quality for for the most commonly assembling dRep clusters from Supp Fig 3. Each bar depicts the n50 (an index of the contiguity of a genome assembly) for the genome from that dRep cluster from a given co-assembly. In almost all cases the genome from the all-fraction co-assembly (bar farthest left) is the highest n50 or close second.

Supplemental Figure 5

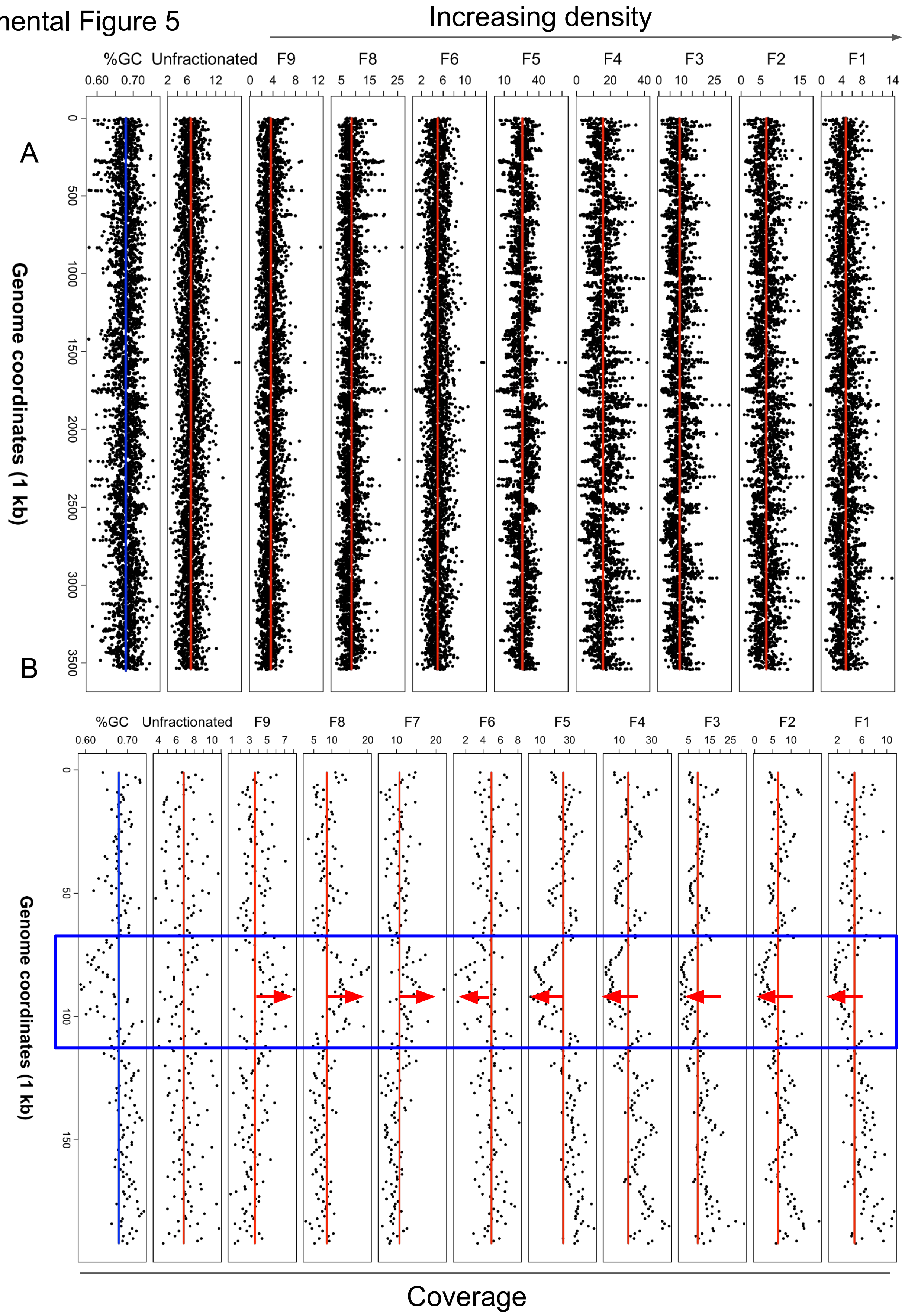


Coverage varies across density gradient for same genome based on GC. A.) coverage and GC for a genome that assembled almost identically across multiple co-assemblies. The vertical axis shows the base-pair position across the sequence. The right-most plot shows the average GC in 1000bp windows. The plots from left to right show the coverage across the sequence in the unfractionated DNA and density fractions from heaviest (F1) to lightest (F9). B.) same plots as in A but for a single large contig. Average coverage in a low-GC region (blue inset in B) varies across density gradient.

Supplemental Figure 6

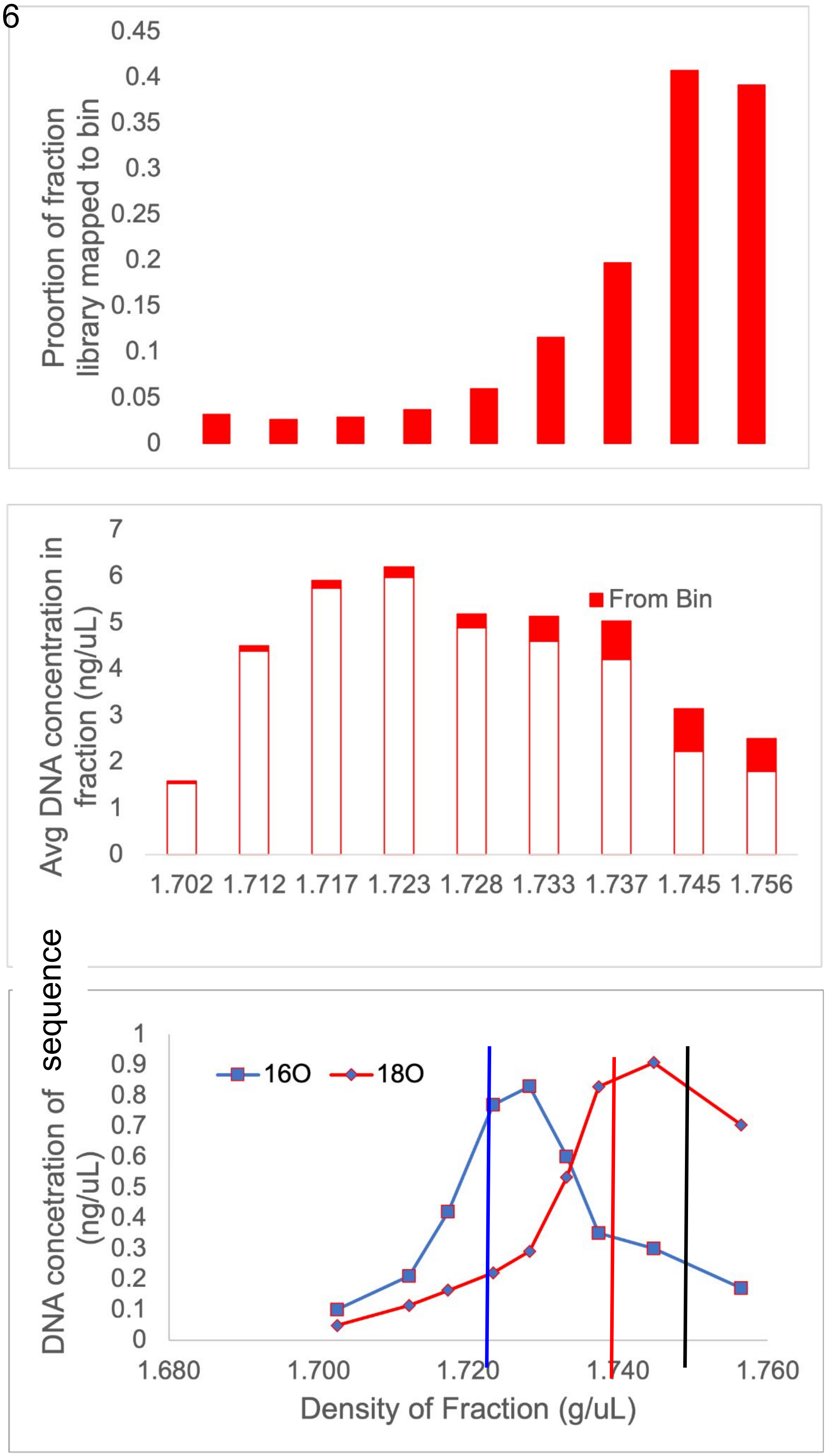


Graphical illustration of AFE calculation for metagenomic sequences using values generated for a theoretical sequence (e.g. contig or MAG). A.) Calculate relative abundance as portion of library mapped to the sequence. B.) Normalize DNA concentration of each sequenced DNA-density gradient fraction by relative abundance of sequence to calculate the DNA concentration of that sequence in each fraction C.) Calculate the average density of that sequence in a given sample weighted by the DNA concentration of the sequence across the density gradient. AFE is the weighted mean density of the sequence in the 18O samples (red vertical line) minus the density in the 16O samples (blue vertical line), as a proportion of the maximum theoretical density shift (black vertical line).

Supplemental Figure 7

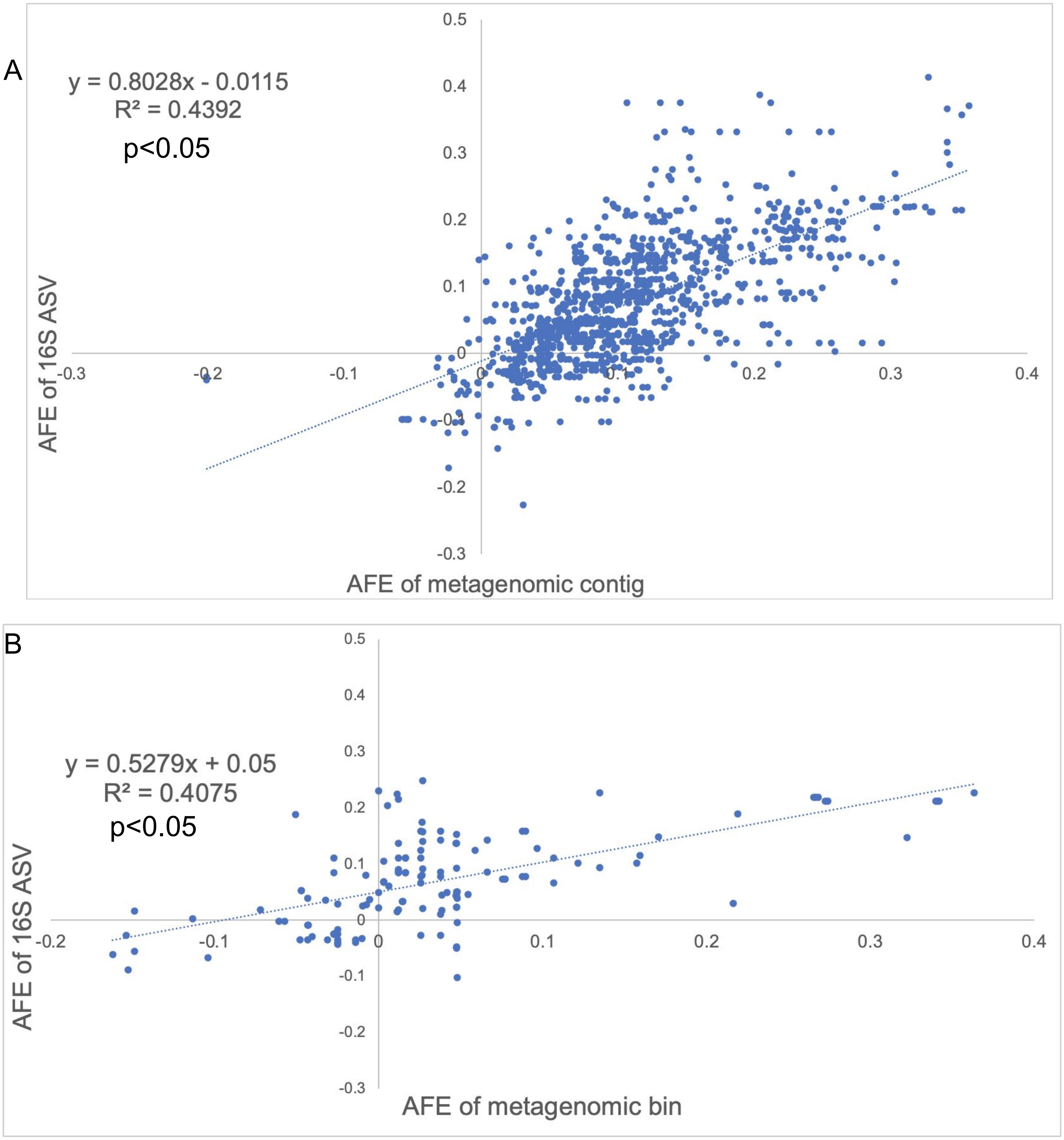


Plots comparing AFE estimated from metagenomic sequences containing 16S rRNA genes and the AFE calculated for ASVs identified from 16S rRNA illumina amplicon sequencing libraries for the same DNA. A.) AFE calculated for metagenomic contigs containing 16S rRNA genes versus AFE estimated for ASVs from 16S amplicons. B.) AFE of the subset of metagenomic bins containing 16S sequences that match 16S-amplicon ASVs.

Supplemental Figure 8

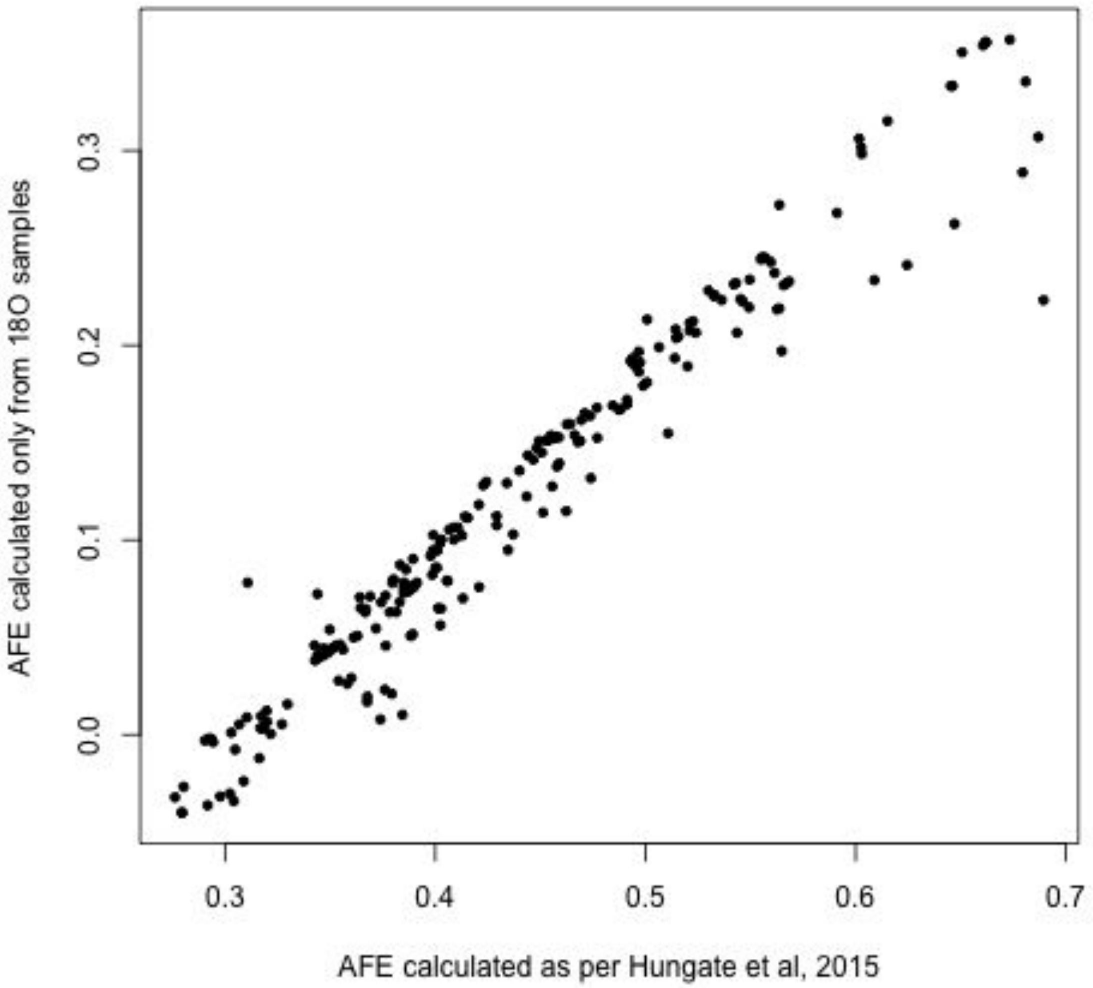


Each point represents a genome assembled from this experiment. The horizontal axis shows the AFE estimated for that genome using the observed weighted mean density of the genome in the 16O samples. The vertical axis shows the AFE estimated for that same genome only using data from the 18O incubations (predicting the natural-abundance WMD of that genome based on GC content).

Supplemental Figure 9

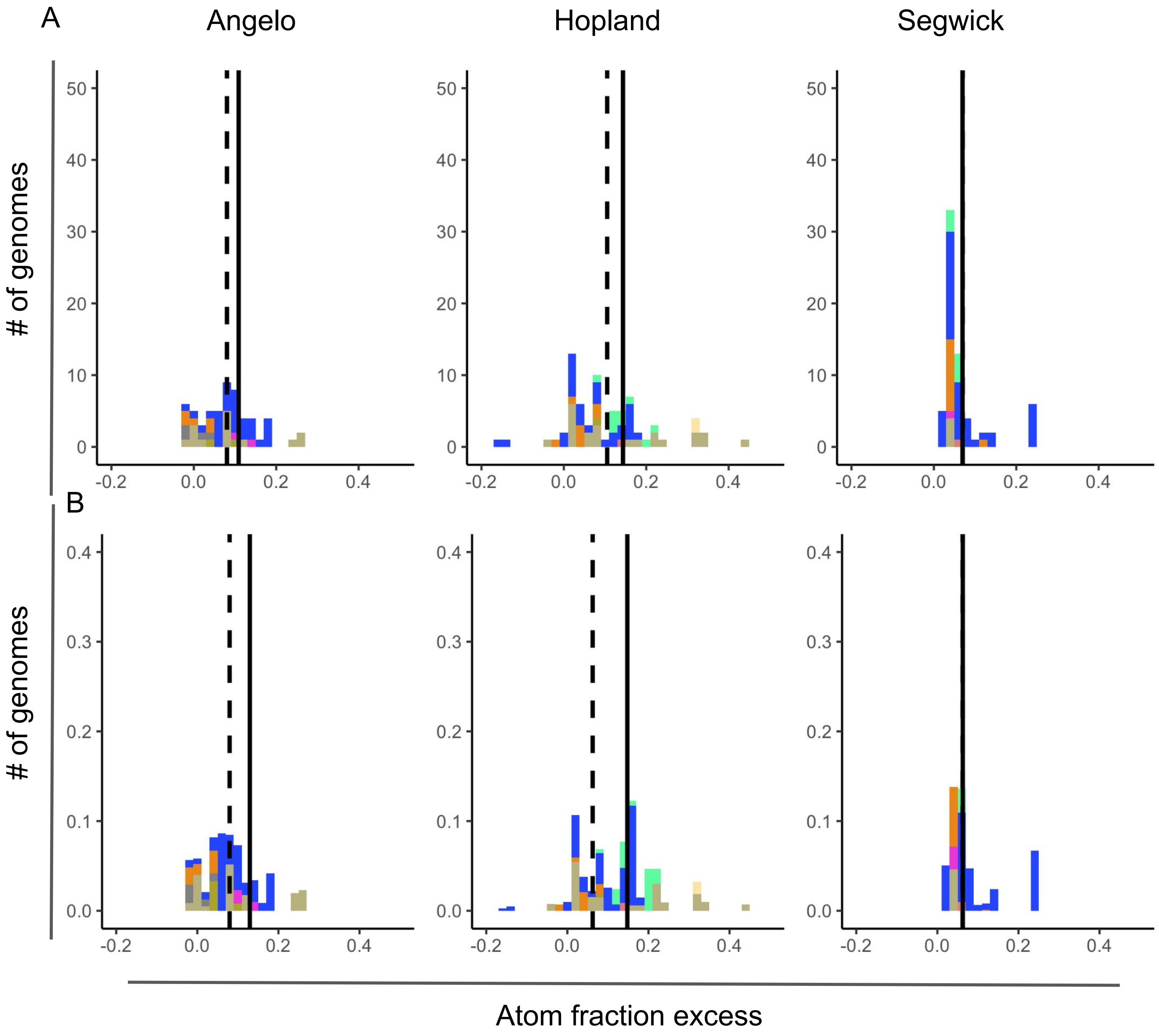


Frequency distribution of AFE values among genomes annotated with the large subunit of aerobic carbon monoxide dehydrogenase (coxL) colored by phylum. The heights of the bars represent count of genomes observed at each AFE level (A) and relative abundance of genomes at each AFE level (B). Solid line represents the average AFE from all bins at that site; dotted line is the average AFE of genomes encoding coxL genes at that site.

Supplemental Figure 10

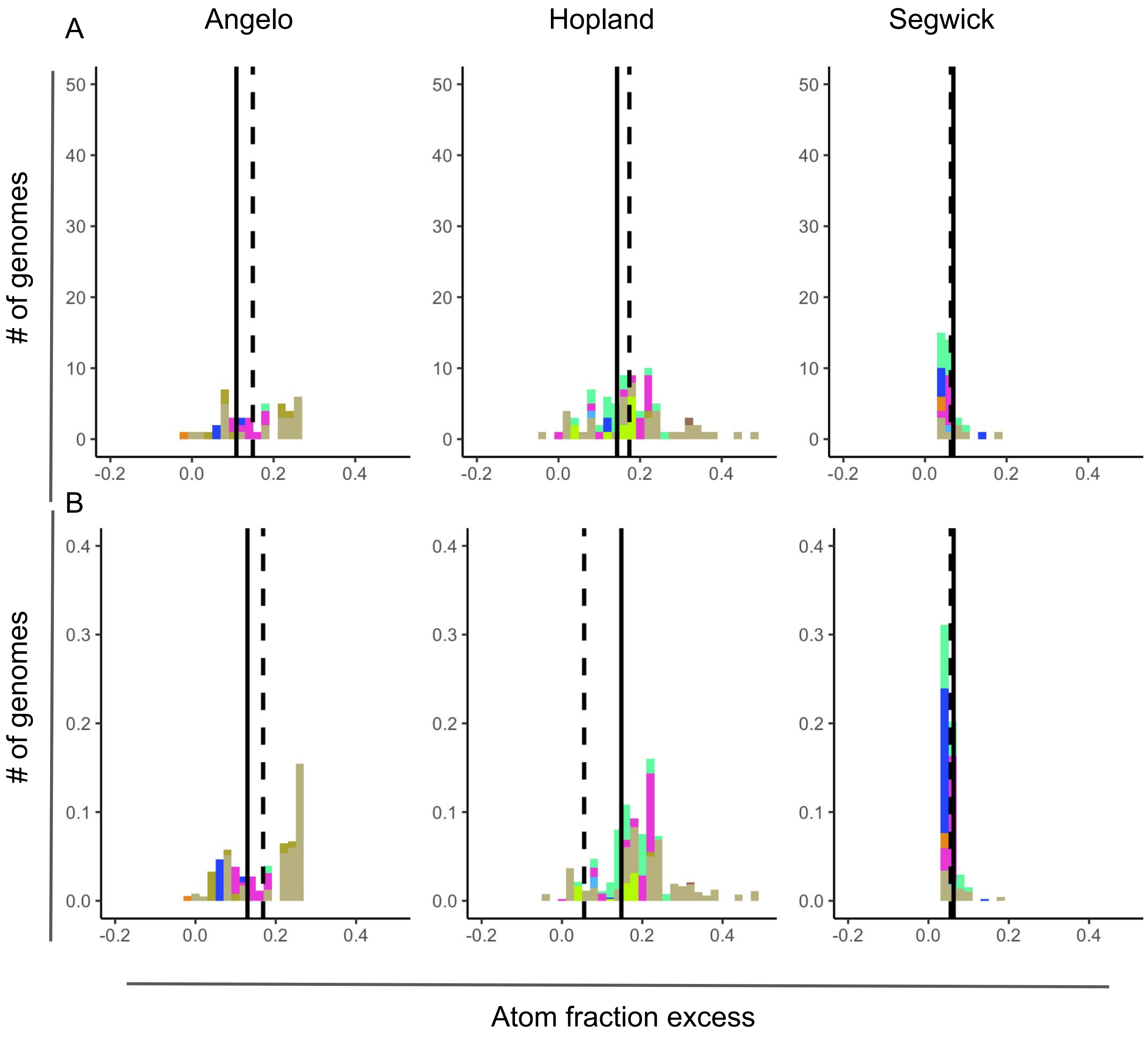


Frequency distribution of AFE values among genomes annotated with methanol dehydrogenase (mxaF) colored by phylum. The heights of the bars represent count of genomes observed at each AFE level (A) and relative abundance of genomes at each AFE level (B). Solid line represents the average AFE from all bins at that site; dotted line is the average AFE of genomes encoding mxaF genes at that site.

Supplemental Figure 11

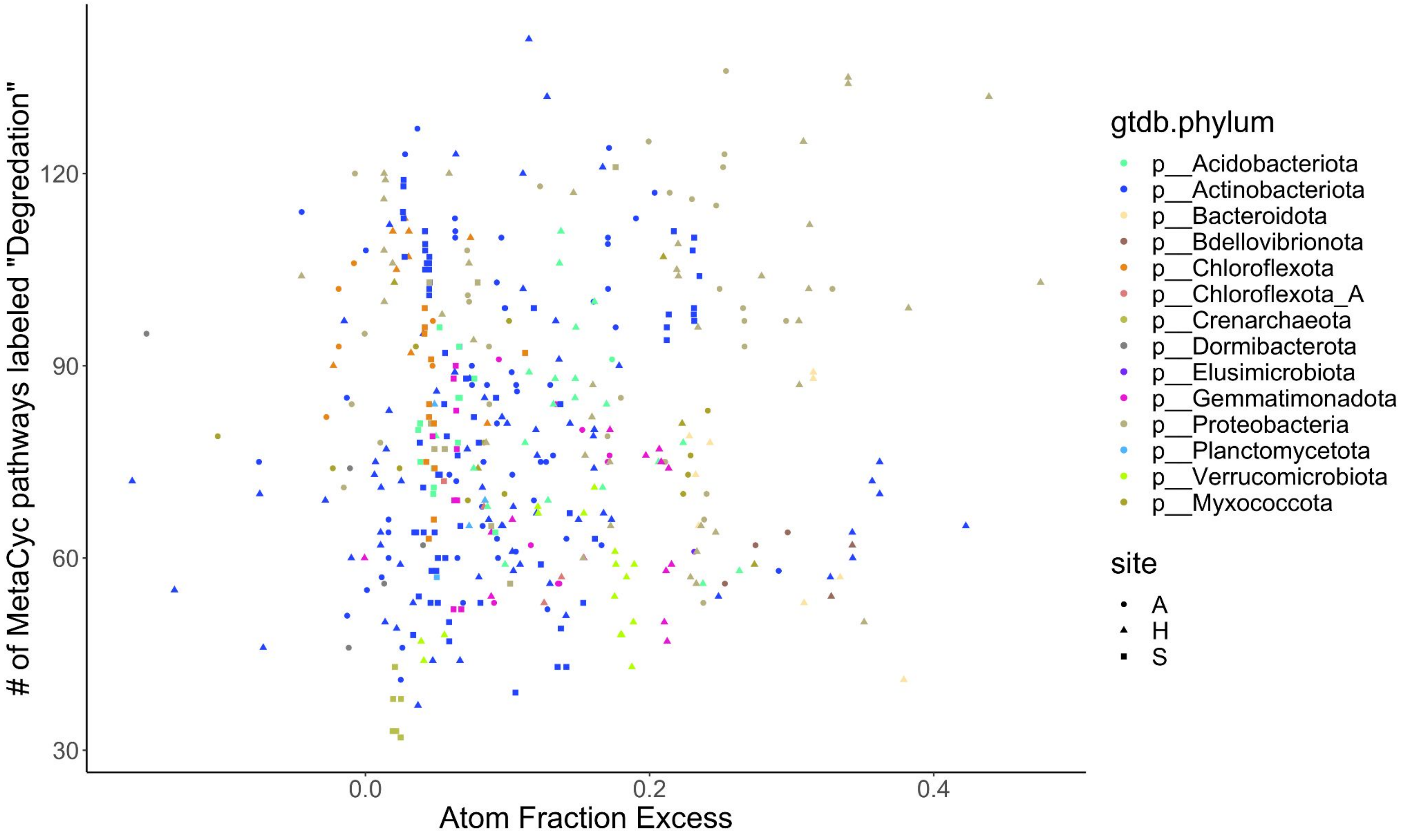


Each point represents a genome. The horizontal axis corresponds to the AFE of that genome, the vertical axis the number of metabolic pathways categorized under degredation from the metaCyc database in that genome. There is no significant correlation between activity measured by AFE and number of metaCyc pathways labeled degradation across sites.

Supplemental Figure 12

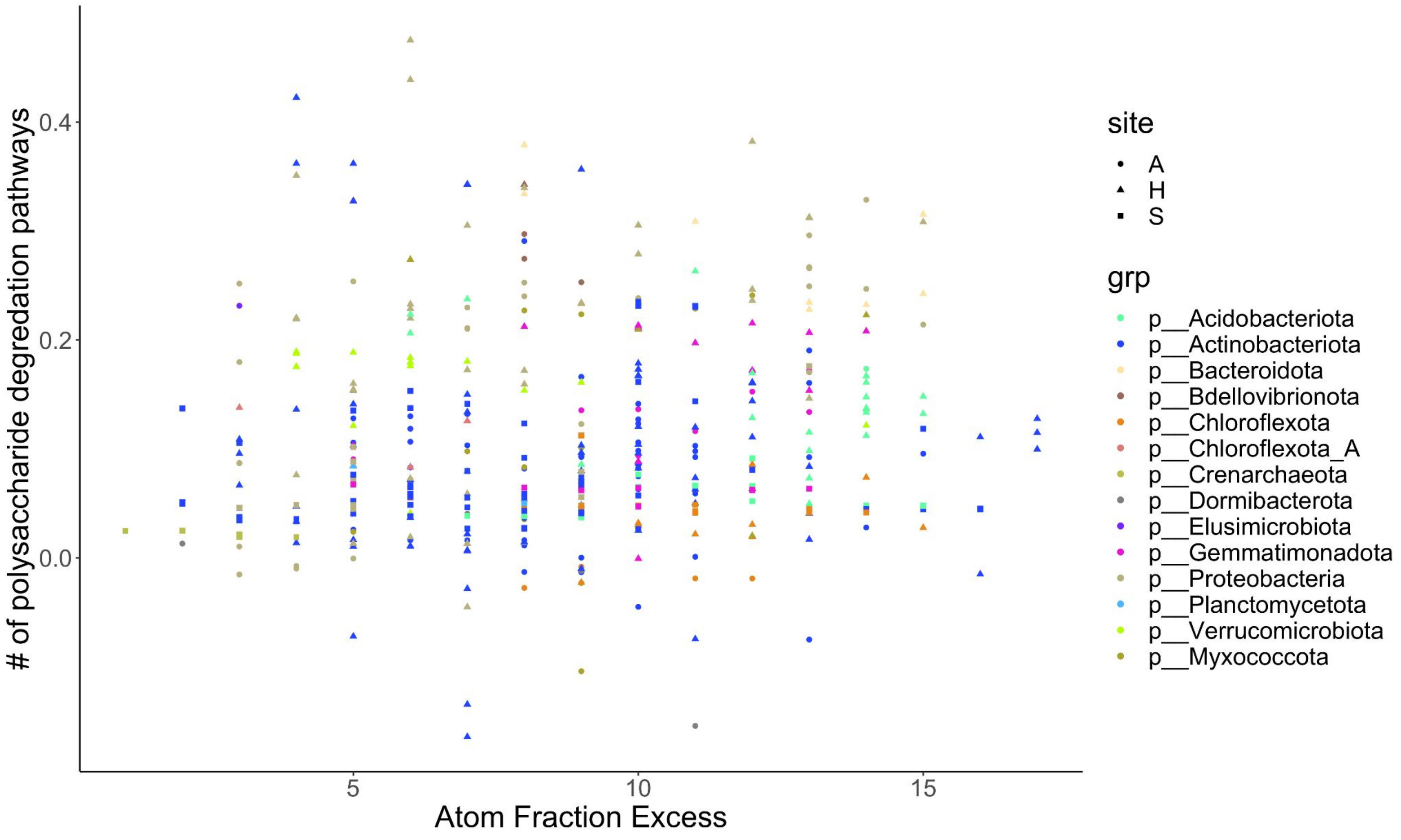


Each point represents a genome. The horizontal axis corresponds to the AFE of that genome, the vertical axis the number of pathways for degradation of polysaccahrides annotated by the DRAM pipeline in that genome. There is no significant correlation between activity measured by AFE and number of polysaccharide degradation pathways annotated in a genome across sites.

Supplemental Figure 13

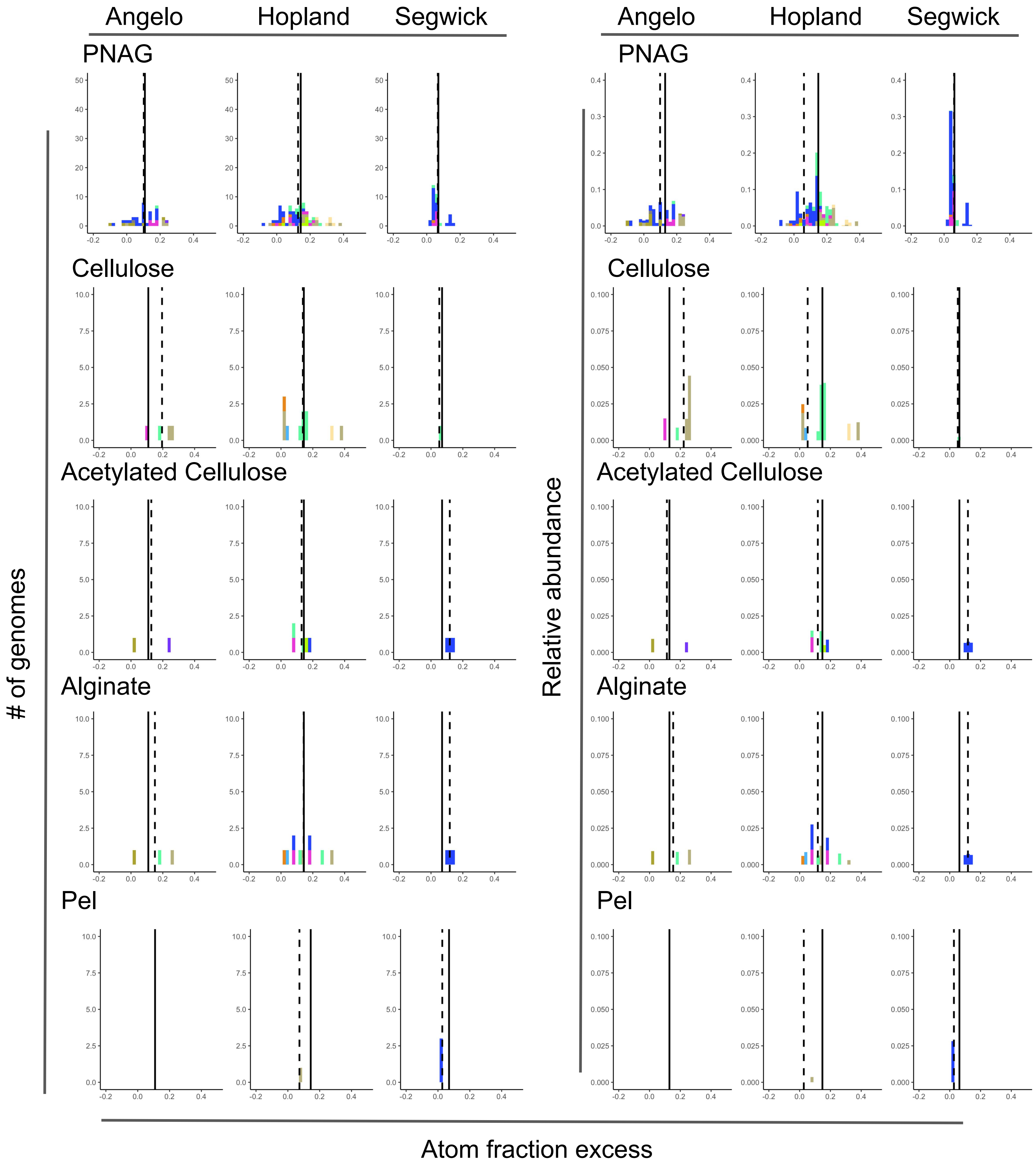


Frequency distribution of AFE values among genomes annotated with biosynthetic gene clusters for synthase-dependent polysaccharides(poly-N-acetylglucosamine (PNAG), cellulose and acetylated cellulose, alginate, and Pel). Plots on the left show counts of genome with each operon and on the left the relative abundance of genomes with each polysaccharide. Solid line represents the average AFE from all bins at that site; dotted line is the average AFE of genomes encoding that polysaccharide biosynthesis pathway at that site.

#### Supplemental Figure 14

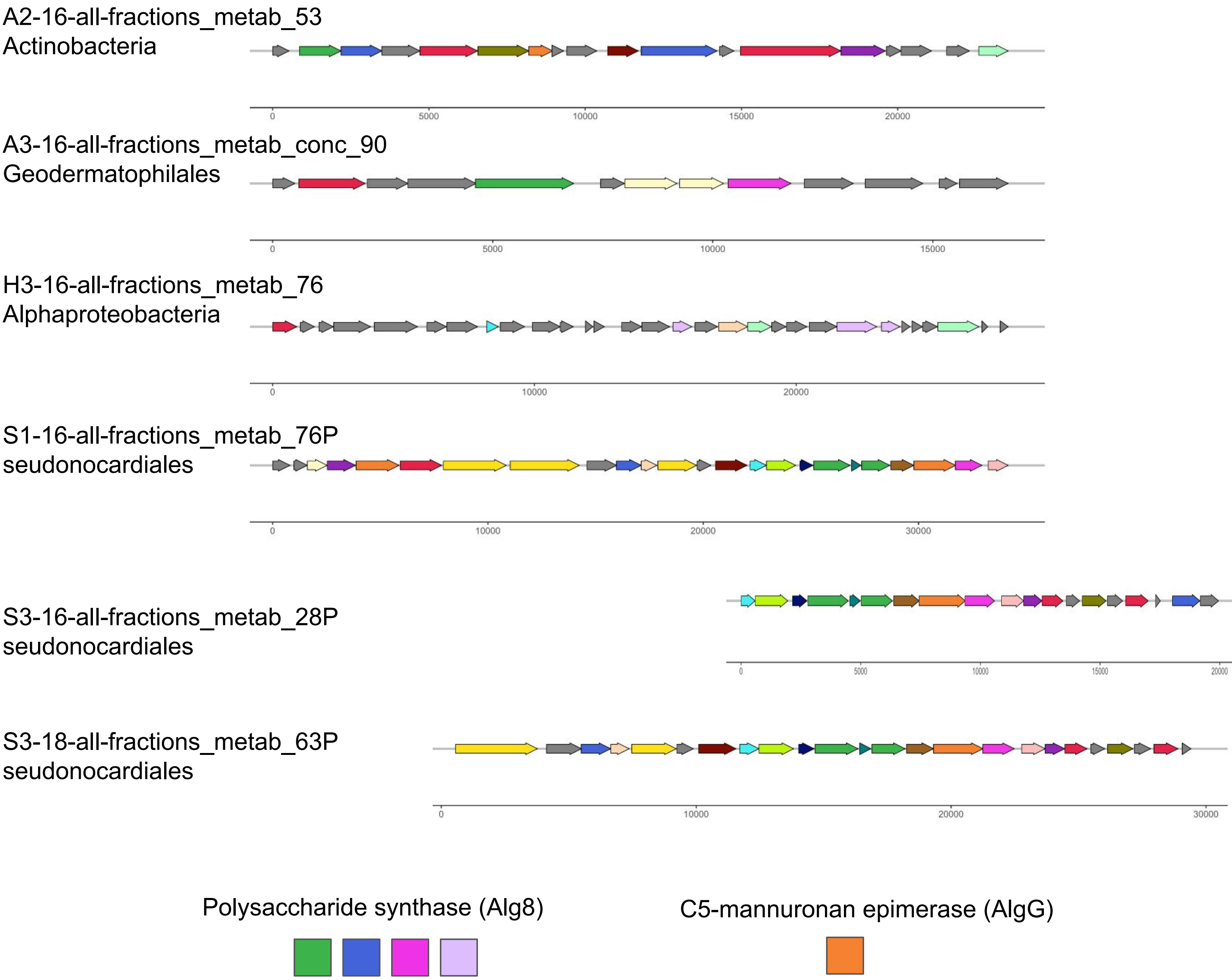


Structure of operons predicted for alginate biosynthesis at Angelo, Hopland and Sedgewick. Genes are colored by membership in protein families generated by clustering all genes in the biosynthetic gene clusters predicted for all five synthase dependent polysaccharide class. Alginate biosynthesis operons from all three genomes at Sedgwick are almost identical.

Supplemental Figure 15

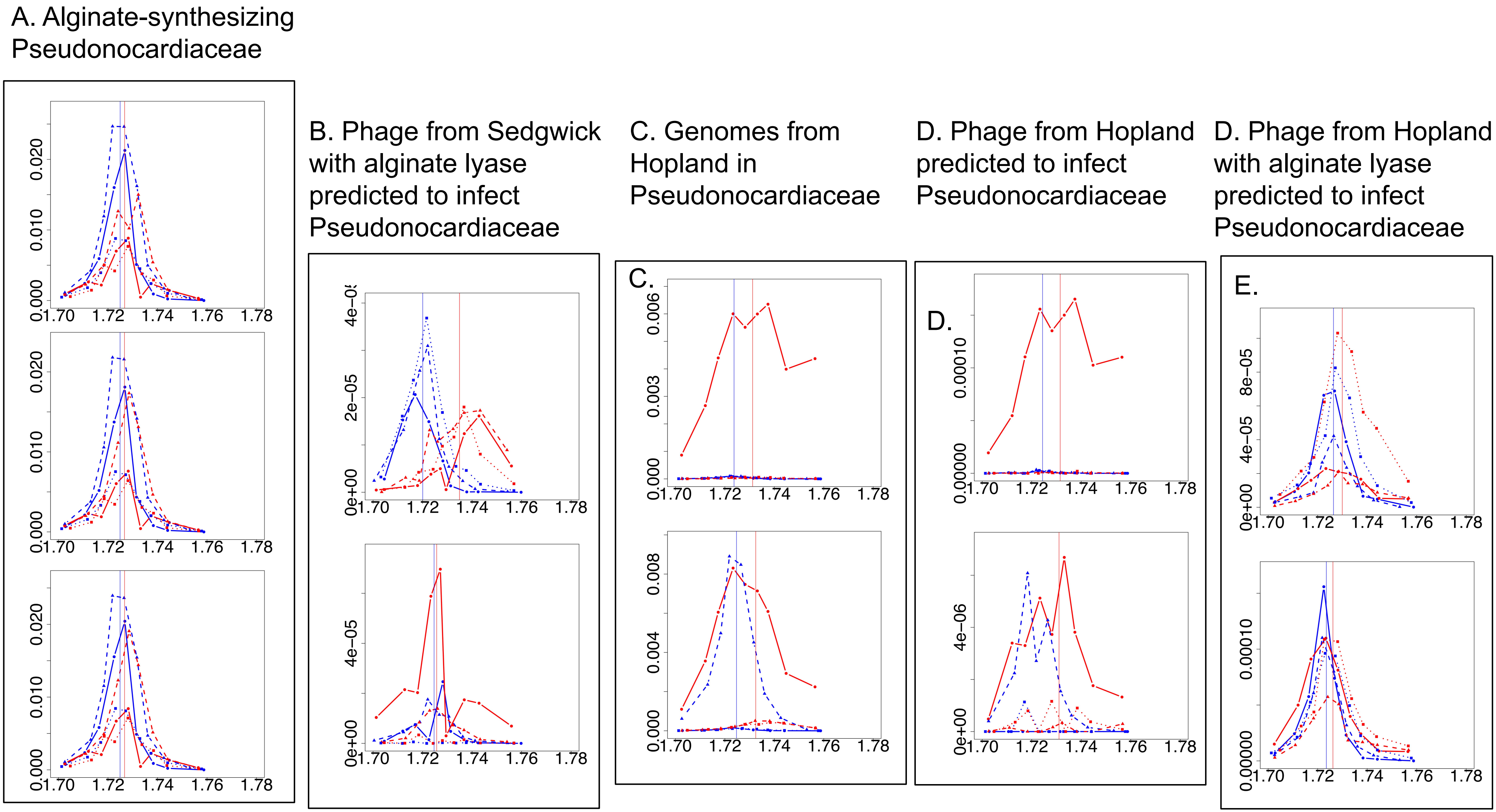


Density distribution of sequences associated with alginate biosynthesis and decomposition at Sedgwick and Hopland. In all plots, blue lines are for natural-abundance 16O treatments and red lines are for 18O treatments. A.) Density distributions of Pseudonocardiaceae (Actinobacteria) genomes predicted to contain alginate biosynthesis operons match the density distributions of phage genomes assembled at Sedgwick predicted to infect actinobacteria in the pseudonocardiales and which were annotated with putative alginate lyase enzymes (B). C.) Genomes from actinobacteria belonging to pseudonocardiales assembled at Hopland have density distributions which match those of phage genomes assembled at Hopland predicted to infect the pseudonocardiales genomes in panel C based on matching CRISPR spacer sequences (D) but not phage contigs assembled at Hopland annotated with alginate lyase and predicted to infect pseudonocardiales.
